# Development of nanobodies as theranostic agents against CMY-2-like class C β-lactamases

**DOI:** 10.1101/2022.07.01.498528

**Authors:** Cawez Frédéric, Paola Sandra Mercuri, Francisco Morales Yanez, Rita Maalouf, Marylène Vandevenne, Frederic Kerff, Virginie Guérin, Jacques Mainil, Damien Thiry, Marc Saulmont, Alain Vanderplasschen, Pierre Lafaye, Gabriel Aymé, Pierre Bogaerts, Mireille Dumoulin, Moreno Galleni

## Abstract

Soluble single-domain fragments derived from the unique variable region of camelid heavy-chain antibodies (VHHs) against enzymes may behave as potent inhibitors. The immunization of alpacas with the CMY-2 β-lactamase led to the isolation of three VHHs that specifically recognized and inhibited CMY-2. The structure of the complex VHH cAb_CMY-2_(254)/CMY-2 was determined by X-ray crystallography. We showed that the epitope is close to the active site and that the CDR3 of the VHH protrudes in the catalytic site. The β-lactamase inhibition was found to follow a mixed profile with a predominant non-competitive component. The three isolated VHHs recognized overlapping epitopes since they behaved as competitive binder. Our study identified a binding site that can be targeted by a new class of β-lactamase’s inhibitors designed with the help of a peptidomimetic approach. Furthermore, the use of mono or bivalent VHH and rabbit polyclonal anti-CMY-2 antibodies enable the development of the first generation of ELISA test for the detection of CMY-2 produced by resistant bacteria.

**IMPORTANCE:** The still increasing antimicrobial resistance in human clinic or veterinary medicine is a major threat for modern chemotherapy. Beside the major caution in the use of current antibiotics, it is important to develop new classes of antibiotics. This work was focused on β-lactamases that are the enzymes involved in the hydrolysis of the major class of antibiotics, the β-lactam compounds. We selected camelid antibodies that inhibit CMY-2, a class C β-lactamase produced by bacteria isolated from the veterinary and human settings. We characterized the conformational epitope present in CMY-2 in order to create a new family of inhibitors based on the paratope of the antibody. Finally, we designed a primary version of a detection system based on an ELISA using VHH and polyclonal antibodies.

## Introduction

Bacterial resistance to antibiotics is unanimously recognized as a major threat in human and veterinary medicine. Nowadays, the antimicrobial resistance counts for 700000 deaths per year in the world, including with 30000 deaths in Europe. This figure could exceed ten million deaths in 2050 if no new treatments and rapid diagnostic assays are developed (1). Among the different classes of antimicrobials, the β-lactam antibiotics are extensively used because of their wide spectrum of action and their low toxicity for the eukaryote cells (2). They are able to specifically neutralize the enzymatic activity of Penicillin Binding Protein (PBP) involved in the formation of the bacterial cell wall (3, 4). Bacteria developed different mechanisms in order to suppress the biological activity of the antibiotics (5–9), the most common one in Gram-negative bacteria being the hydrolysis of the β-lactam ring by the expression of enzymes called β-lactamases.

Up to date, more than 2800 bla genes are known (10). Their related enzymes are categorized into 4 molecular classes A, B, C and D based exclusively on their primary sequence (11). Another classification of the β-lactamases in different functional sub-groups is based on both molecular and functional characteristics (12).

The misuse and the intensive use of antibiotics lead to the selection of multidrug-resistant (MDR) bacteria which is unaffected by the presence antibiotics belonging to at least three different classes (13). Therefore, it is essential to develop new diagnostic assays in order to faster detect the presence of β-lactamase(s) that favor(s) the rapid implementation of infection control measures and circumvent the nosocomial infections. In addition, it is essential to develop new inhibitors able to block the β-lactamase activity by targeting binding sites that are not tolerant to mutations.

To develop new inhibitors, one strategy consists to select inhibitory antibodies that serve to the development of new β-lactamase inhibitors by peptidomimetics (14). VHH, also referred as nanobody, is the single-domain fragment corresponding to the binding domain of camelid heavy-chain antibodies (HCAbs), constitutes a potential candidate in view to obtain inhibitory antibodies against the β-lactamases. They are exclusively found in camelids (VHH) or in cartilaginous fish (V-NAR) (15). Despite their small size (15kDa), they are able to interact with their antigen with a high affinity and specificity (16). In addition, VHHs present unique properties including an easy recombinant production in *E. coli*, an easiness to modify the properties of the nanobody by protein engineering. Moreover, VHHs are able to inhibit activity of some enzymes as previously described for the lysozyme (17) and for the β-lactamases TEM-1 (18) and VIM-4 (19).

The “RUBLA” project which studied the distribution of *bla* genes in bovine *E. coli* isolates in Wallonia, Belgium highlighted that the bla_CMY-2_ coding for the cephalosporinase CMY-2 is characterized by the broadest geographic spread (20). In addition, *Enterobacteriaceae* strains expressing this β-lactamase were isolated from animal (22) and human sources (23).

Phenotypic assays were developed to detect the production of AmpC by testing the susceptibility of the strains for ceftazidime, cefoxitin and cefepime. In addition, phenotypic confirmation tests imply the use of inhibitors such as cloxacillin or boronic acid derivatives (24). Nevertheless, those methods cannot identify the different sub-classes of AmpC. The assays must be complemented by molecular approaches such as PCR or micro-array when available (25). Those methods are expensive, time-consuming and generally used only for reference laboratories and research settings (26). Furthermore, expression of multiple β- lactamases (27) or other resistance features such as the decrease of porins expression could result in more complex susceptibility patterns (28). Altogether, those observations clearly demonstrate the real necessity to develop new diagnostic approaches for the veterinary and the clinic in order to detect AmpC easily and with a high specificity.

On the other hand, treatments generally employed to treat infections of *Enterobacteriaceae* expressing AmpC consist in the use of carbapenems, cefepime or tazobactam in association with piperacillin (29). Newer β-lactamase inhibitors as avibactam or vaborbactam have also a high potency against AmpC activity (30). However, it is expected to favor the apparitions of resistance against those antibiotics and inhibitors, specifically against carbapenems (31).

In this work, we developed nanobodies (VHHs) in order to set-up a sandwich ELISA for the detection of CMY-2 and to find inhibitors able to neutralize the β-lactamase activity. We first isolated eight VHHs, belonging to three families, that recognized CMY-2. The results highlighted a high specificity for their antigen but rapid dissociation rates of the complexes VHHs/CMY-2. We could stabilize the complex with the development of a homo-bivalent VHH. Competition assays demonstrated also an overlapping epitope of the VHHs for CMY-2. With the help of an ELISA test, we could detect the production of CMY-2-like enzymes in a collection of human and veterinary bacterial isolates. The second goal was to follow the effect of the VHHs binding on the CMY-2 activity. We found that the VHHs behaved as non-competitive inhibitors of CMY-2 and that the nature of the substrate affected the inhibition patterns of the nanobodies. The crystallographic structure of the complex formed by CMY-2 and the VHH cAb_CMY-2_ (254) allowed to define the epitope recognized by the VHH, the nature of the paratope and to confirm the inhibitory mechanisms highlighted by the kinetic studies. Altogether, those results provide new insights for diagnostic and inhibitory antibodies development against class C β-lactamases.

## Results

Construction of an immune VHHs library and selection of CMY-2 targeting binders. From the blood of an alpaca (*Vicugna pacos*) immunized with CMY-2 -lactamase, an immune library was constructed. Three rounds of panning using this library were performed to enrich the library in phage particles exposing CMY-2-specific VHHs using established protocols (47). Ninety clones of each round of panning, randomly selected, were screened by indirect ELISA to detect the presence of VHHs specific for CMY-2. Eight clones gave a positive signal in the ELISA and their gene was sequenced. The results of the sequencing indicated 3 genetically different VHHs (cAb_CMY-2_(250), cAb_CMY-2_(254) and cAb_CMY-2_(272)) based on the sequence of the Complementarity Determining Regions (CDRs) (Fig. 1). CDR2 and CDR3 of the VHHs cAb_CMY-2_(250) and cAb_CMY-2_(272) present a deletion of one and four amino acids, respectively, compared to the VHH cAb_CMY-2_(254). The FR4 sequences are identical for the three VHH but we observed a deletion of two amino acids in the FR3 of the VHH cAb_CMY-2_(272). Finally, despite if the main mutations are identified into the CDRs, the VHH cAb_CMY-2_(254) is also characterized by additional mutations in the frameworks regions. Based on these results, VHHs cAb_CMY-2_(250), cAb_CMY-2_(254) and cAb_CMY-2_(272) were produced and purified to complete their analysis.

**FIG 1.**
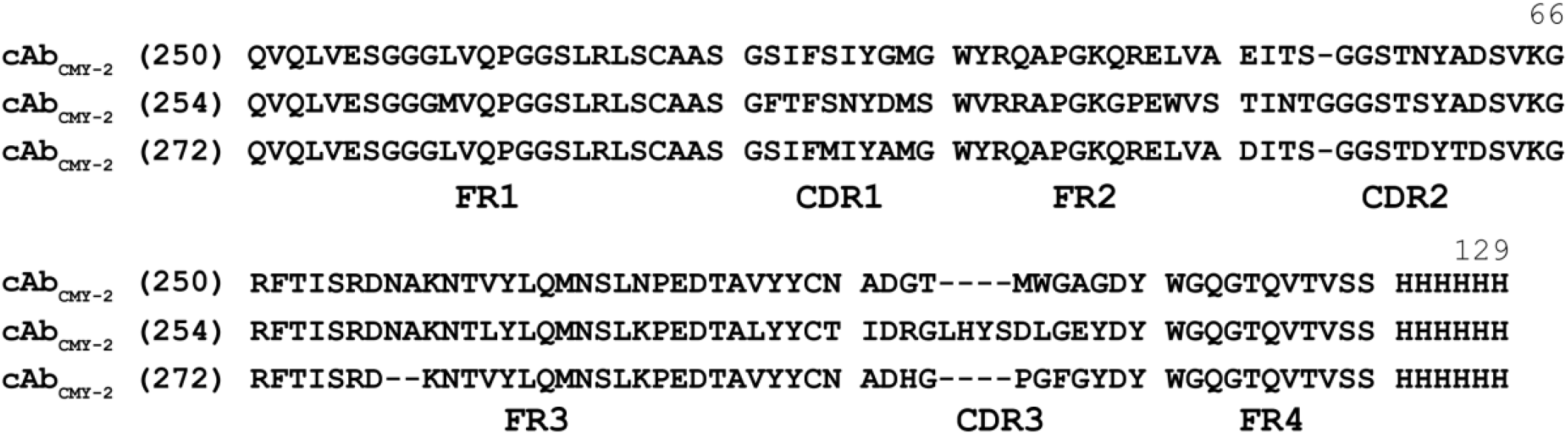
Sequence alignment of VHHs directed against CMY-2. FR: framework, CDR: complementarity determining region.

Binding characterization of the VHHs cAb_CMY-2_(250), cAb_CMY-2_(254) and cAb_CMY-2_(272) by bio-layer interferometry (BLI). In order to determine the specificity of the VHHs cAb_CMY-2_(250), cAb_CMY-2_(254) and cAb_CMY-2_(272), qualitative binding measurements were performed to assess their ability to interact with different representatives of all the molecular classes of β-lactamases (BLAs). The three VHHs did not recognize β-lactamases from classes A, B and D (Fig. 2A-C). Remarkably, the VHHs cAb_CMY-2_(254) and cAb_CMY-2_(272) interacted only with CMY-2-like enzymes (Fig. 2B & 2C) but not with others class C β-lactamases indicating their remarkable specificity. On the contrary, cAb_CMY-2_(250) displayed a cross reaction with the AmpC P99 of *Enterobacter cloacae* (Fig. 2A). Quantitative binding measurements were performed in order to measure the kinetic (k_on_, k_off_) and equilibrium (K_D_) constants (Fig. 2D-F). The association kinetic constants (k_on_) of the three VHHs against CMY-2 ranged from 10^5^ to 10^6^ M^-1^s^-1^ highlighting a rapid association of the VHHs to their target (TABLE 1). Nevertheless, all complexes were quite unstable given their relatively fast dissociation kinetic (k_off_ > 5 10^-3^ s^-1^) that leads to overall moderate affinities of VHHs for CMY-2 (K_D_ > 60 nM). The comparison of binding properties of the three VHHs showed that cAb_CMY-2_(254) presents a dissociation that is up to 20 times lower than the two others VHHs (Fig. 2B) indicating this VHH is able to form a more stable complex than cAb_CMY-2_ (250) and cAb_CMY-2_ (272).

**TABLE 1.**
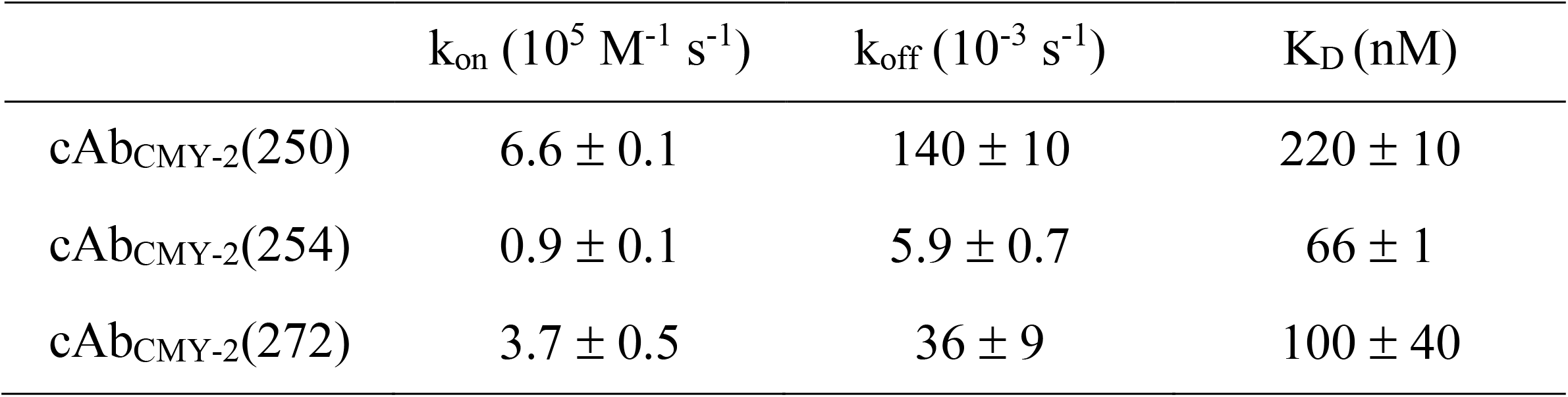
Kinetic (k_on_, k_off_) and equilibrium (K_D_) constants determined by BLI. k_on_ and k_off_ values obtained were derived from a global fit of at least seven analyte concentrations (i.e. with 7 analyte concentrations illustrated on sensorgrams in Fig. 2) with the equation of a 1:1 binding model. All values described in the table above include averages and standard deviations calculated from two independent experiments.

**FIG 2.**
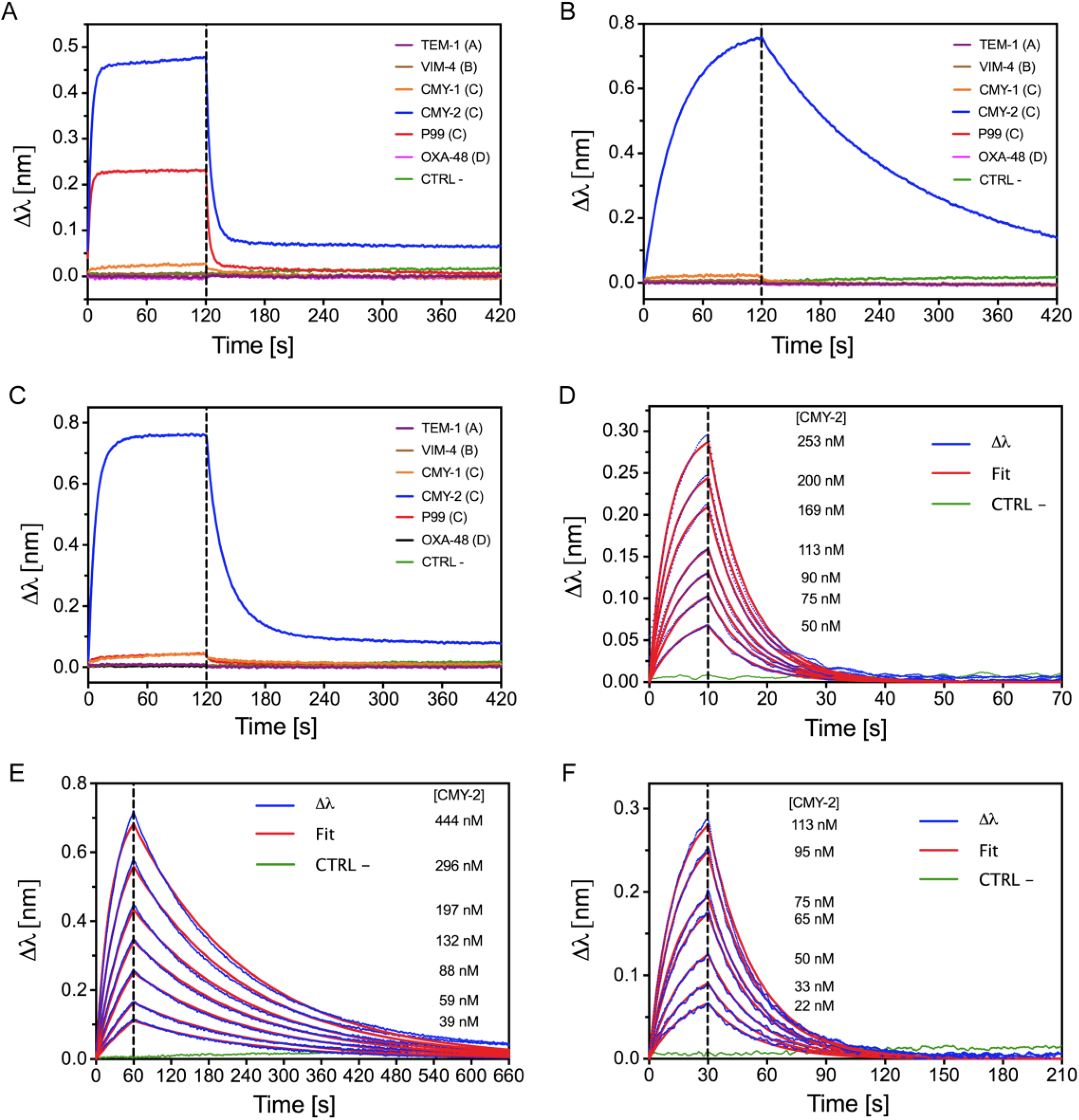
Binding characterization of the three selected VHHs by bio-layer interferometry. Qualitative binding specificity of (A) cAb_CMY-2_(250), (B) cAb_CMY-2_(254) and (C) cAb_CMY-2_(272), respectively. Names and classes (in brackets) of the tested β-lactamases are indicated on the figure. Quantitative binding measurements of (D) cAb_CMY-2_(250), (E) cAb_CMY-2_(254) and (F) cAb_CMY-2_(272), respectively. The experimental data (Δλ, blue) recorded with seven different concentrations were fitted using a 1:1 binding model (red). The negative control (CTRL -, green) corresponds to CMY-2 directly loaded on the bio-sensor. Those experiments were carried out twice independently.

Competition binding assay of VHHs directed against CMY-2. We determined if the three nanobodies could recognize the same or different epitopes. Therefore, we carried out competition binding assays by BLI based on a premix method. This latter consists in (i) binding a biotinylated VHH on a streptavidin bio-sensor (SA sensor) and (ii) measuring its association rate in presence of increasing molar ratios of soluble complexes formed by a second VHH and CMY-2. The three possible combinations of complexes (cAb_CMY-2_(250)/CMY-2, cAb_CMY-2_(254)/CMY-2 and cAb_CMY-2_(272)/CMY-2) were assessed for each biotinylated VHH. A decrease in the binding rate of the VHH cAb_CMY-2_ (254) for increased molar ratios of all VHH/CMY-2 complexes was observed (Fig. 3), indicating that the VHH cAb_CMY-2_(254) epitope is, at least, partially overlapping with the epitopes of the two others VHHs. Those results were confirmed when either VHHs cAb_CMY-2_(250) and cAb_CMY-2_(272) were immobilized on the bio-sensor confirming the overlap of the epitopes of the three VHHs (Fig. 1S).

**FIG 3.**
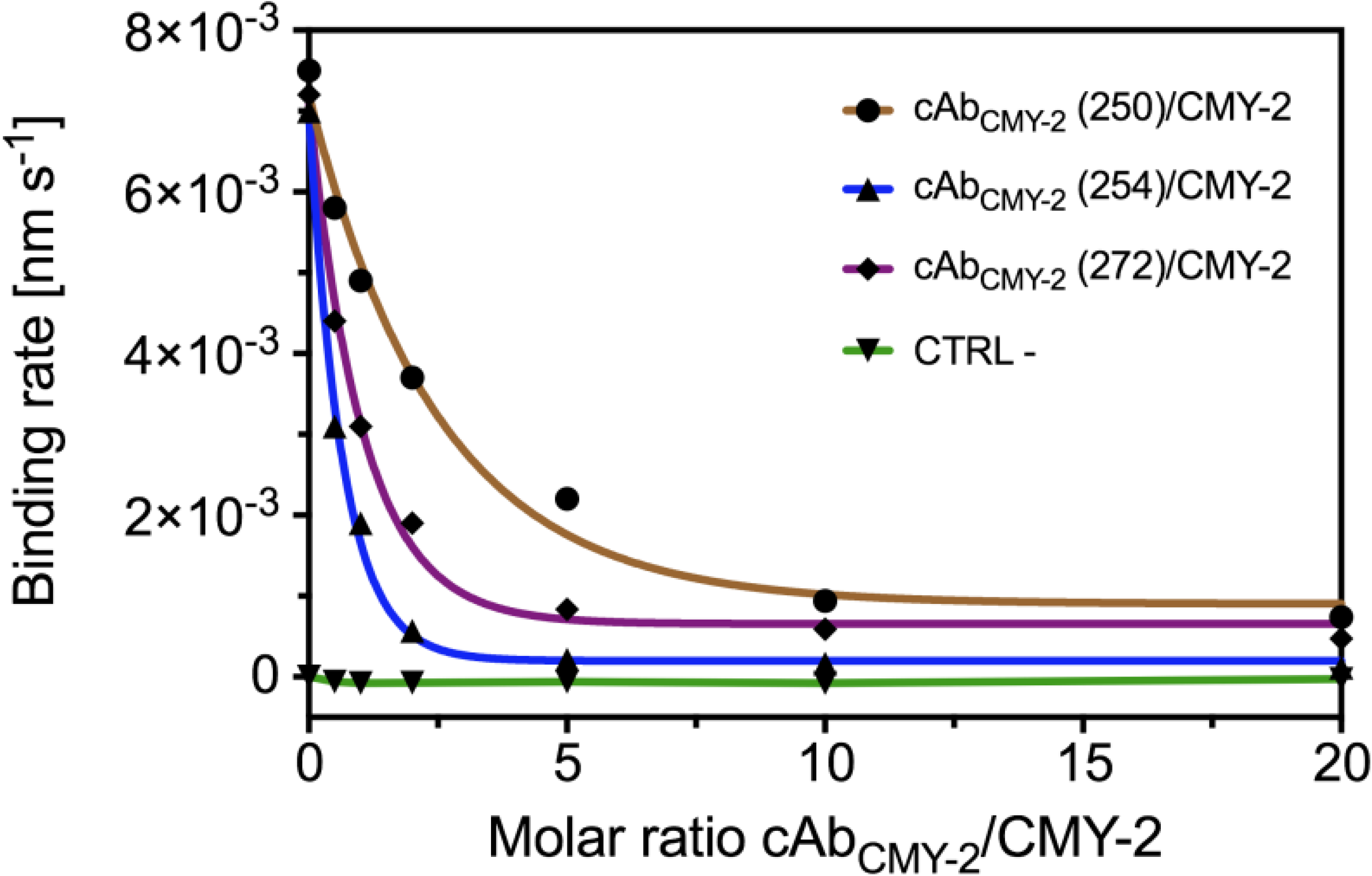
Competition binding assay between VHHs directed against CMY-2 monitored by BLI. The biotinylated VHH cAb_CMY-2_(254) was loaded on a streptavidine bio-sensor (SA sensor) while the analyte corresponds to complexes cAb_CMY-2_(250)/CMY-2 (brown), cAb_CMY-2_(254)/CMY-2 (blue) and cAb_CMY-2_(272)/CMY-2 (purple) in different molar ratios. The negative control (CTRL -, green) corresponds to the signal recorded when the complex cAb_CMY-2_(254)/CMY-2 was directly loaded on the non-functionalized bio-sensor. All the binding rates were calculated by fitting a simple exponential mathematic model to the first 120 seconds of the association phase. This experiment was realized twice independently.

Binding properties of rabbit polyclonal antibodies directed against CMY-2 (pAbs). The epitope overlapping of the three VHHs does not allow the development of a VHH sandwich ELISA. To circumvent this issue, we produced and characterized rabbit polyclonal antibodies directed against CMY-2 (anti-CMY-2 pAbs) with the purpose to be used in pair with cAb_CMY-2_(254), the VHH forming the more stable complex with CMY-2. Firstly, the specificity of anti-CMY-2 pAbs was analyzed by indirect ELISA where a panel of β-lactamases representative of all classes representing were coated on a 96-well plate (Fig. 4A). The data clearly indicated that pAbs directed against CMY-2 were unspecific since they recognized different members of class C β-lactamases such as CMY-1 and P99. A second assay via BLI was also conducted to further investigate the specificity of pAbs. Biotinylated β-lactamases were immobilized on streptavidin bio-sensors and then, sensors were immerged into a pAbs solution (Fig. 4B). This experiment confirms the lack of specificity since bindings were also measured for P99 and CMY-2. However, no interaction was detected for CMY-1 demonstrating the lack of specificity can depend on the assay setting and/or the type of immobilization. Finally, the dissociation constant (k_off_) was evaluated for pAbs by BLI (Fig. 4C) which, as commonly observed for polyvalent antibodies, presented a high avidity characterized by a slow dissociation phase (k_off_ = 3.6 ± 0.9 10^-5^ s^-1^) (32).

**FIG 4.**
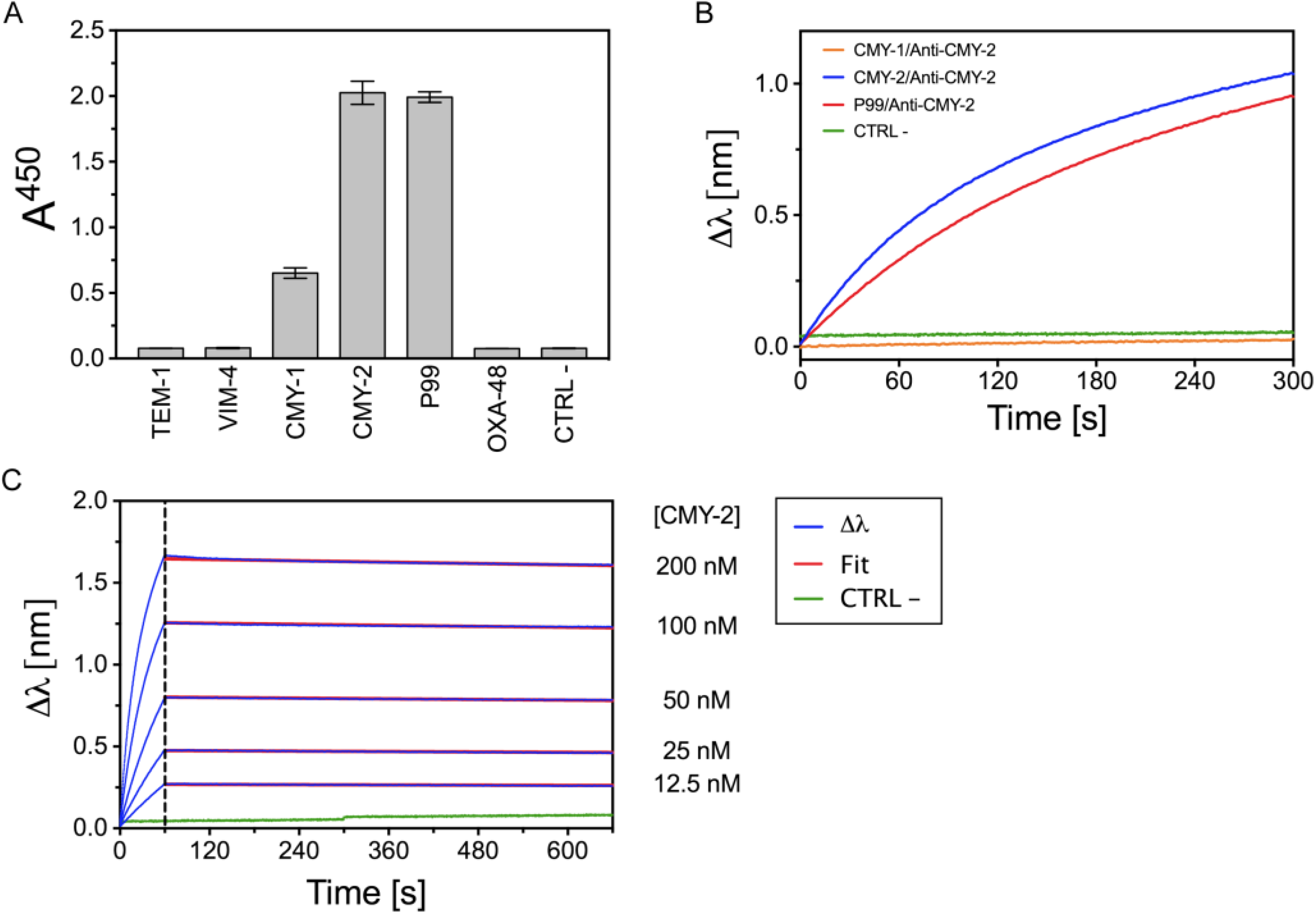
Binding properties of rabbit polyclonal antibodies directed against the β-lactamase CMY-2 (anti-CMY-2 pAbs). (A) Indirect ELISA where the antigens were absorbed on a maxisorp plate (except for the CTRL -) to investigate the specificity of pAbs. Values correspond to means and standard deviations from duplicates. (B) Qualitative binding specificity assay of pAbs for CMY-2 monitored by BLI. (C) Quantitative binding measurements of pAbs for CMY-2 realized by BLI. The experimental data (Δλ, blue) recorded with five different concentrations were fitted using a 1:1 binding model (red). The negative control (CTRL -, green) corresponds to anti-CMY-2 pAbs directly loaded on the bio-sensor. The BLI experiments were realized twice independently.

Sandwich ELISA for the detection of the β-lactamase CMY-2. The first step consisted to evaluate the limits of detection (LOD) for a sandwich ELISA by using cAb_CMY-2_(254) as capture antibody and pAbs for the detection (full blue line) or, inversely, the pAbs as capture antibody and the VHH for the detection (Fig. 5A). The LOD values were calculated from an average Abs^450^ of the CTRL (–) plus three times the standard deviation and corresponded to 13.3 and 3.9 ng/mL using the VHH cAb_CMY-2_(254) as capture and detection antibodies, respectively. Compared to LOD values reported in the literature (LOD = 0.86 ng/mL) (33), our ELISA is characterized by high LOD values that are mainly due to the fast dissociation of VHH/CMY-2 complexes. This phenomenon has a negative impact on the limit of detection of our antigens. Moreover, the use of pAbs as capture antibody and VHH for the revelation did not bring any gain since the responses (measured as Abs^450^) were lower compared to the initial settings. On the other hand, the specificity of the sandwich ELISA assay was checked on purified enzymes (Fig. 5B). Both assays clearly allow the specific detection of CMY-2 since no signal was measured for others families of β-lactamases.

**FIG 5.**
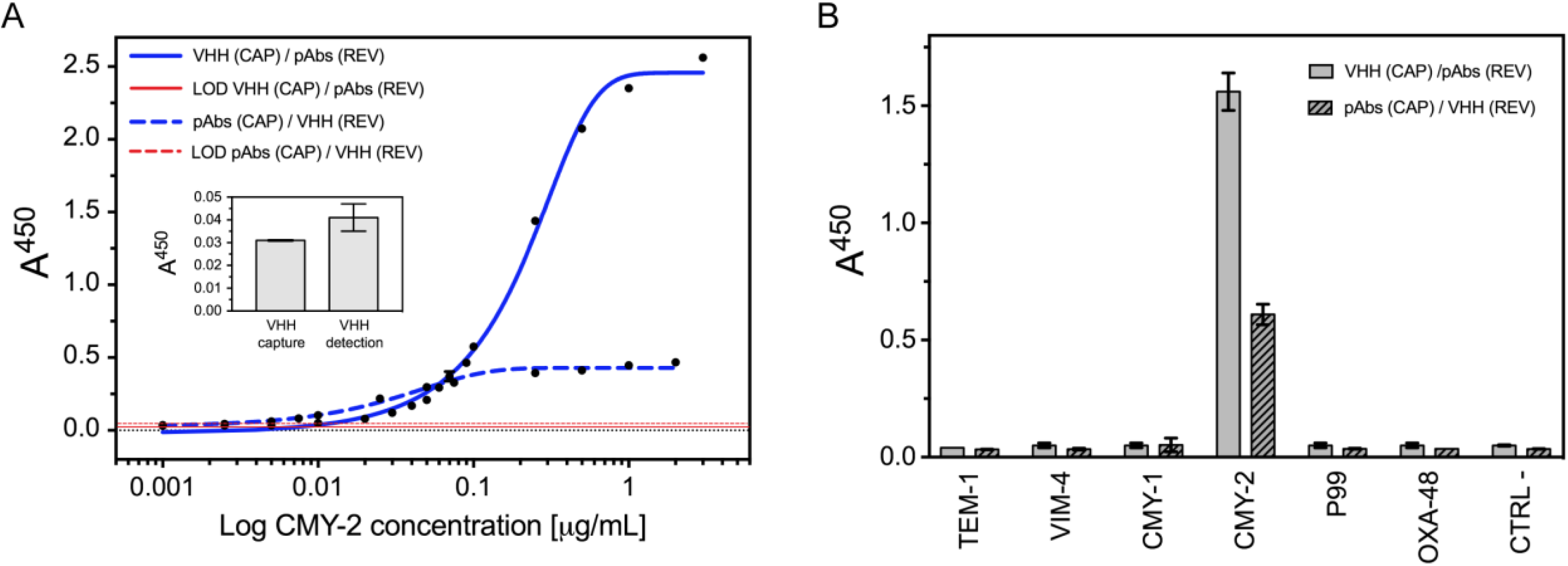
Sandwich ELISA on purified enzymes for CMY-2 detection. (A) Limits of detection (LOD) where VHH cAb_CMY-2_(254) was employed as antibody for capture and pAbs for detection (blue full line) or inversely (blue dotted line). Curves were fitted with equation I (TEXT. S1). The inset includes the Abs^450^ for the negative control (CTRL -). The LOD was calculated from an average Abs^450^ of the CTRL (–) plus three times the standard deviation. They are represented in red full and dotted lines for VHH used as capture and detection antibody, respectively. (B) Specificity assessment using the VHH cAb_CMY-2_(254) as capture agent and the pAbs for revelation (grey) or inversely (grey, pattern). In both experiments, the CTRL - corresponds to the same ELISA without antigen. All averages and standard deviations are results from at least twice measurements. CAP: capture, REV: revelation.

Production and characterization of the bivalent VHH cAb_CMY-2_(254)_BIV_. In order to increase the sensitivity of the ELISA assay, we designed a bivalent VHH based on the VHH cAb_CMY-2_(254). This genetically engineered bivalent antibody (cAb_CMY-2_(254)_BIV_) consists in the fusion of two VHHs cAb_CMY-2_(254) in tandem repeats, joined by a peptide linker (GGGS)_3_(34). The cAb_CMY-2_(254)_BIV_ was produced and purified in comparable amount than the monovalent VHH. In addition, no degradation of the tandem was observed. Bivalent VHHs may exhibit an increased apparent affinity (or avidity) due to a significant decrease in the dissociation rate constant (decreased k_off_ value) from the immobilized antigen (35). As expected, the dissociation rate significantly decreased for the VHH cAb_CMY-2_(254)_BIV_ (k_off_ = 3.8 ± 0.4 x 10^-4^ s^-1^) compared to its monovalent counterpart (k_off_ = 6.3 ± 0.5 x 10^-3^ s^-1^) (Fig. 6). This implies a more stable Antigen/Antibody complex.

**FIG 6.**
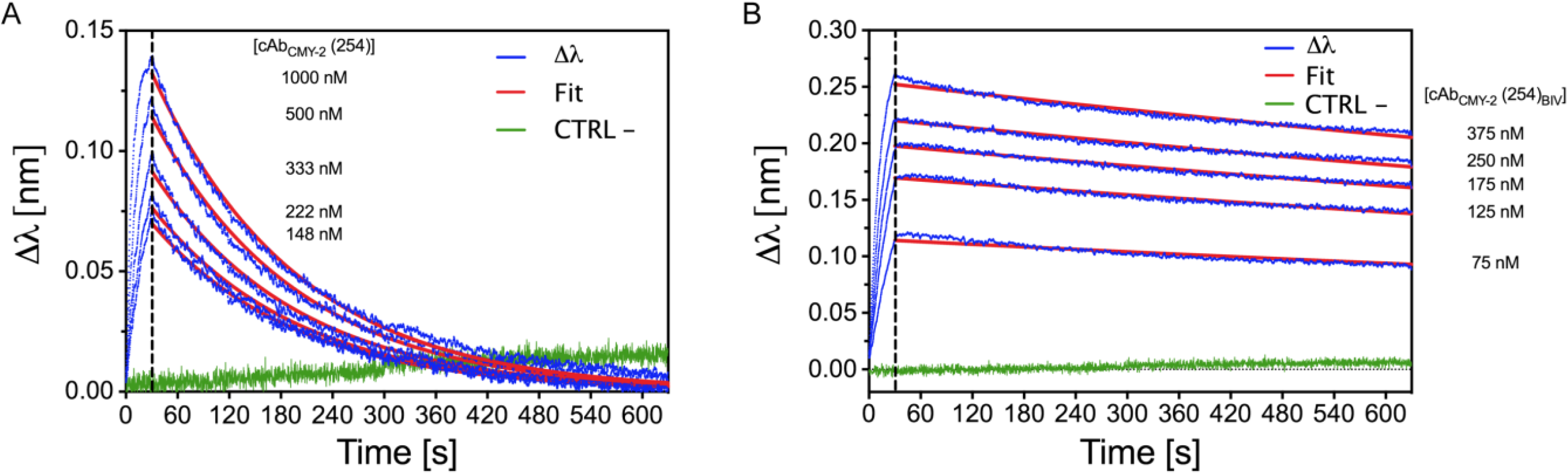
Quantitative binding measurements of the monovalent VHH cAb_CMY-2_(254) (A) and the bivalent VHH cAb_CMY-2_(254)_BIV_ (B) performed by BLI. The experimental data (Δλ, blue) recorded with five different concentrations were fitted using a 1:1 binding model (red). The negative control (CTRL -, green) corresponds to VHHs directly loaded on the bio-sensor. Experiments were realized twice independently.

Sandwich ELISA for the detection of CMY-2 using the bivalent VHH cAb_CMY-2_(254)_BIV_. As for the monovalent counterpart, we evaluated the limits of detection (LOD) for a sandwich ELISA by using the bivalent VHH cAb_CMY-2_(254)_BIV_ as capture antibody and pAbs for the detection (full blue line) and, in parallel, the pAbs as capture antibody and the bivalent VHH for the detection (dotted blue line) (Fig. 7A). Those sets up provided LOD values around 2.3 and 1.4 ng/mL using the VHH cAb_CMY-2_(254)_BIV_ as capture and detection antibodies, respectively. The use of the bivalent VHH clearly improves the detection of CMY-2. Moreover, those configurations seem to not impede the specificity of the ELISA system (Fig. 7B). Indeed, higher sensibility of the assay combined with its high specificity can allow the use of the bivalent VHH cAb_CMY-2_ (254)_BIV_ for the detection of CMY-2 produced in bacterial isolates.

**FIG 7.**
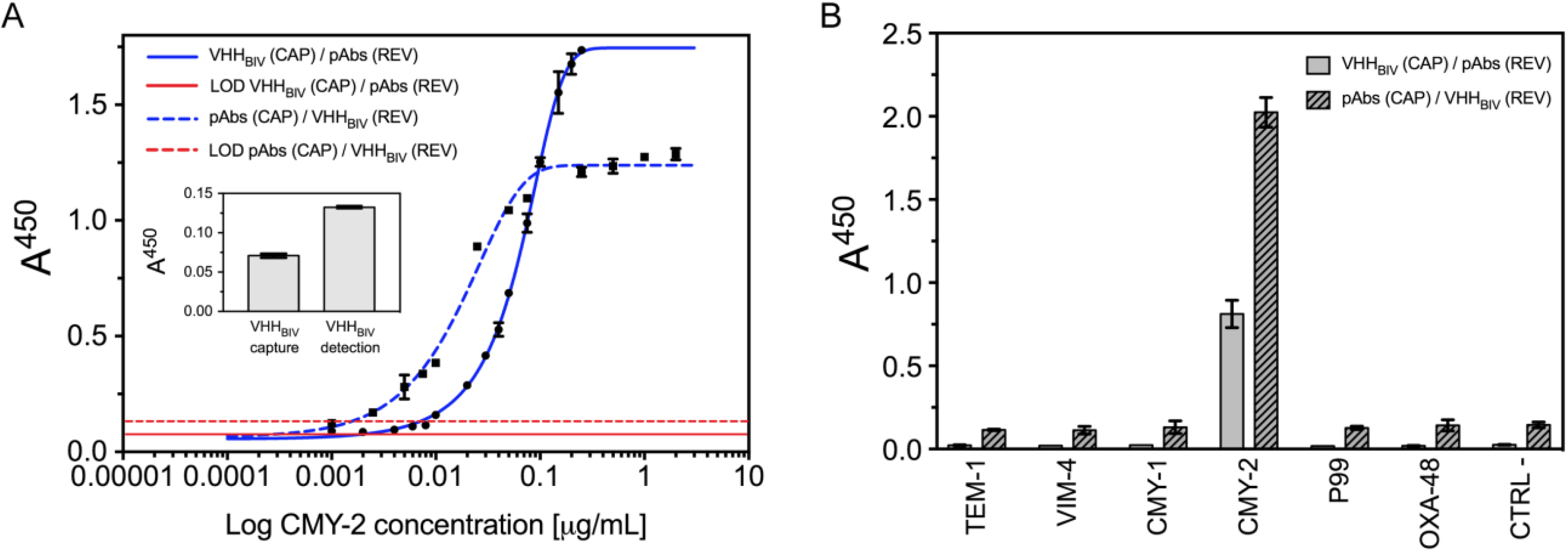
Sandwich ELISA for CMY-2 detection using the bivalent VHH cAb_CMY-2_(254)_BIV_ (A) Limits of detection where the VHH cAb_CMY-2_(254)_BIV_ was used as antibody for capture and pAbs for detection (blue full line) or inversely (blue dotted line). The inset includes the Abs^450^ for the negative control (CTRL -). The LOD was calculated from an average Abs^450^ of the CTRL (–) plus three times the standard deviation. They are represented in red full and dotted lines for VHH used as capture and detection antibody, respectively. (B) Specificity measurement by use of the VHH cAb_CMY-2_(254)_BIV_ as antibody for capture and the pAbs for the detection (grey) or inversely (pattern, grey). The negative control (CTRL -) corresponds to the ELISA without antigen. All averages and standard deviations are results from at least twice measurements. CAP: capture, REV: revelation.

### Detection of CMY-2 β-lactamase in bovine and human bacterial isolates

According to the previous results for purified CMY-2, three sandwich ELISAs were designed for the direct detection of CMY-2 produced in bovine and human bacterial isolates. In these experiments, the monovalent VHH cAb_CMY-2_(254) was only used as capture antibody and anti-CMY-2 pAbs for the detection. At the opposite, the bivalent VHH was used as antibody for capture and revelation since both configurations improved the sensitivity of the assay. Foremost, the three different ELISA allowed the detection of a large panel of CMY-2 sub-group variants such as CMY-16 and CMY-60 (TABLE 2). In addition, no cross reactions were observed for other class C β-lactamases such as the CMY-10 (CMY-1-like), DHA, ACT or subclasses of AmpC expressed in bovine isolates. Interestingly, the bivalent VHH cAb_CMY-2_(254)_BIV_ used as antibody for the capture and the revelation ensured a higher sensitivity for the detection of β-lactamases belonging to the CMY-2 sub-group compared to the monovalent VHH cAb_CMY-2_(254) used as capture antibody. Effectively, only 17 on 22 bacterial isolates presenting one gene coding for CMY-2 sub-group variant were detected via the monovalent VHH versus 21 isolates for the bivalent VHH cAb_CMY-2_(254)_BIV_ (TABLE 2). Our data suggest that our sandwich ELISA can be employed to detect specifically β-lactamases from the CMY-2 sub-group with the interest to use the bivalent VHH cAb_CMY-2_(254)_BIV_ in order to increase the sensitivity of detection.

**TABLE 2.**
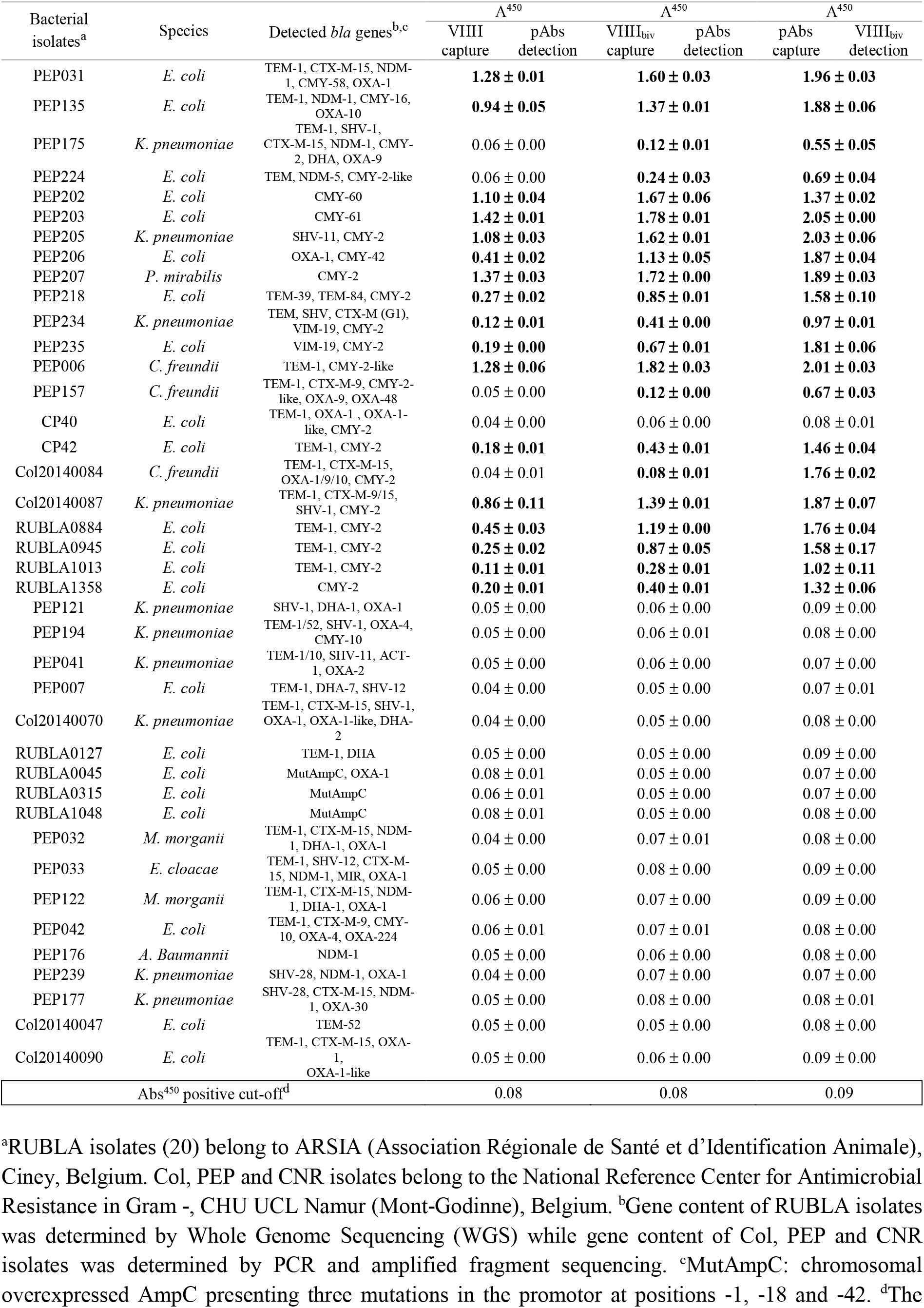
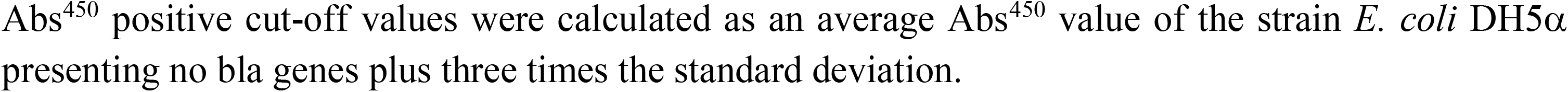
Detection of CMY-2 sub-group β-lactamases in bovine and human bacterial isolates by a sandwich ELISA. All positive strains in the ELISA are indicated in bold.

| Bacterial isolates<sup>a</sup> | Species | Detected *bla* genes<sup>b,c</sup> | A<sup>450</sup> |  | A<sup>450</sup> |  | A<sup>450</sup> |  |
| --- | --- | --- | --- | --- | --- | --- | --- | --- |
|  |  |  | VHH capture | pAbs detection | VHH<sub>biv</sub> capture | pAbs detection | pAbs capture | VHH<sub>biv</sub> detection |
| PEP031 | *E. coli* | TEM-1, CTX-M-15, NDM-1, CMY-58, OXA-1 | **1.28 ± 0.01** |  | **1.60 ± 0.03** |  | **1.96 ± 0.03** |  |
| PEP135 | *E. coli* | TEM-1, NDM-1, CMY-16, OXA-10 | **0.94 ± 0.05** |  | **1.37 ± 0.01** |  | **1.88 ± 0.06** |  |
| PEP175 | *K. pneumoniae* | TEM-1, SHV-1, CTX-M-15, NDM-1, CMY-2, DHA, OXA-9 | 0.06 ± 0.00 |  | **0.12 ± 0.01** |  | **0.55 ± 0.05** |  |
| PEP224 | *E. coli* | TEM, NDM-5, CMY-2-like | 0.06 ± 0.00 |  | **0.24 ± 0.03** |  | **0.69 ± 0.04** |  |
| PEP202 | *E. coli* | CMY-60 | **1.10 ± 0.04** |  | **1.67 ± 0.06** |  | **1.37 ± 0.02** |  |
| PEP203 | *E. coli* | CMY-61 | **1.42 ± 0.01** |  | **1.78 ± 0.01** |  | **2.05 ± 0.00** |  |
| PEP205 | *K. pneumoniae* | SHV-11, CMY-2 | **1.08 ± 0.03** |  | **1.62 ± 0.01** |  | **2.03 ± 0.06** |  |
| PEP206 | *E. coli* | OXA-1, CMY-42 | **0.41 ± 0.02** |  | **1.13 ± 0.05** |  | **1.87 ± 0.04** |  |
| PEP207 | *P. mirabilis* | CMY-2 | **1.37 ± 0.03** |  | **1.72 ± 0.00** |  | **1.89 ± 0.03** |  |
| PEP218 | *E. coli* | TEM-39, TEM-84, CMY-2 | **0.27 ± 0.02** |  | **0.85 ± 0.01** |  | **1.58 ± 0.10** |  |
| PEP234 | *K. pneumoniae* | TEM, SHV, CTX-M (G1), VIM-19, CMY-2 | **0.12 ± 0.01** |  | **0.41 ± 0.00** |  | **0.97 ± 0.01** |  |
| PEP235 | *E. coli* | VIM-19, CMY-2 | **0.19 ± 0.00** |  | **0.67 ± 0.01** |  | **1.81 ± 0.06** |  |
| PEP006 | *C. freundii* | TEM-1, CMY-2-like | **1.28 ± 0.06** |  | **1.82 ± 0.03** |  | **2.01 ± 0.03** |  |
| PEP157 | *C. freundii* | TEM-1, CTX-M-9, CMY-2-like, OXA-9, OXA-48 | 0.05 ± 0.00 |  | **0.12 ± 0.00** |  | **0.67 ± 0.03** |  |
| CP40 | *E. coli* | TEM-1, OXA-1, OXA-1-like, CMY-2 | 0.04 ± 0.00 |  | 0.06 ± 0.00 |  | 0.08 ± 0.01 |  |
| CP42 | *E. coli* | TEM-1, CMY-2 | **0.18 ± 0.01** |  | **0.43 ± 0.01** |  | **1.46 ± 0.04** |  |
| Col20140084 | *C. freundii* | TEM-1, CTX-M-15, OXA-1/9/10, CMY-2 | 0.04 ± 0.01 |  | **0.08 ± 0.01** |  | **1.76 ± 0.02** |  |
| Col20140087 | *K. pneumoniae* | TEM-1, CTX-M-9/15, SHV-1, CMY-2 | **0.86 ± 0.11** |  | **1.39 ± 0.01** |  | **1.87 ± 0.07** |  |
| RUBLA0884 | *E. coli* | TEM-1, CMY-2 | **0.45 ± 0.03** |  | **1.19 ± 0.00** |  | **1.76 ± 0.04** |  |
| RUBLA0945 | *E. coli* | TEM-1, CMY-2 | **0.25 ± 0.02** |  | **0.87 ± 0.05** |  | **1.58 ± 0.17** |  |
| RUBLA1013 | *E. coli* | TEM-1, CMY-2 | **0.11 ± 0.01** |  | **0.28 ± 0.01** |  | **1.02 ± 0.11** |  |
| RUBLA1358 | *E. coli* | CMY-2 | **0.20 ± 0.01** |  | **0.40 ± 0.01** |  | **1.32 ± 0.06** |  |
| PEP121 | *K. pneumoniae* | SHV-1, DHA-1, OXA-1 | 0.05 ± 0.00 |  | 0.06 ± 0.00 |  | 0.09 ± 0.00 |  |
| PEP194 | *K. pneumoniae* | TEM-1/52, SHV-1, OXA-4, CMY-10 | 0.05 ± 0.00 |  | 0.06 ± 0.01 |  | 0.08 ± 0.00 |  |
| PEP041 | *K. pneumoniae* | TEM-1/10, SHV-11, ACT-1, OXA-2 | 0.05 ± 0.00 |  | 0.06 ± 0.00 |  | 0.07 ± 0.00 |  |
| PEP007 | *E. coli* | TEM-1, DHA-7, SHV-12 | 0.04 ± 0.00 |  | 0.05 ± 0.00 |  | 0.07 ± 0.01 |  |
| Col20140070 | *K. pneumoniae* | TEM-1, CTX-M-15, SHV-1, OXA-1, OXA-1-like, DHA-2 | 0.04 ± 0.00 |  | 0.05 ± 0.00 |  | 0.08 ± 0.00 |  |
| RUBLA0127 | *E. coli* | TEM-1, DHA | 0.05 ± 0.00 |  | 0.05 ± 0.00 |  | 0.09 ± 0.00 |  |
| RUBLA0045 | *E. coli* | MutAmpC, OXA-1 | 0.08 ± 0.01 |  | 0.05 ± 0.00 |  | 0.07 ± 0.00 |  |
| RUBLA0315 | *E. coli* | MutAmpC | 0.06 ± 0.01 |  | 0.05 ± 0.00 |  | 0.07 ± 0.00 |  |
| RUBLA1048 | *E. coli* | MutAmpC | 0.08 ± 0.01 |  | 0.05 ± 0.00 |  | 0.08 ± 0.00 |  |
| PEP032 | *M. morganii* | TEM-1, CTX-M-15, NDM-1, DHA-1, OXA-1 | 0.04 ± 0.00 |  | 0.07 ± 0.01 |  | 0.08 ± 0.00 |  |
| PEP033 | *E. cloacae* | TEM-1, SHV-12, CTX-M-15, NDM-1, MIR, OXA-1 | 0.05 ± 0.00 |  | 0.08 ± 0.00 |  | 0.09 ± 0.00 |  |
| PEP122 | *M. morganii* | TEM-1, CTX-M-15, NDM-1, DHA-1, OXA-1 | 0.06 ± 0.00 |  | 0.07 ± 0.00 |  | 0.09 ± 0.00 |  |
| PEP042 | *E. coli* | TEM-1, CTX-M-9, CMY-10, OXA-4, OXA-224 | 0.06 ± 0.01 |  | 0.07 ± 0.01 |  | 0.08 ± 0.00 |  |
| PEP176 | *A. Baumannii* | NDM-1 | 0.05 ± 0.00 |  | 0.06 ± 0.00 |  | 0.08 ± 0.00 |  |
| PEP239 | *K. pneumoniae* | SHV-28, NDM-1, OXA-1 | 0.04 ± 0.00 |  | 0.07 ± 0.00 |  | 0.07 ± 0.00 |  |
| PEP177 | *K. pneumoniae* | SHV-28, CTX-M-15, NDM-1, OXA-30 | 0.05 ± 0.00 |  | 0.08 ± 0.00 |  | 0.08 ± 0.01 |  |
| Col20140047 | *E. coli* | TEM-52 | 0.05 ± 0.00 |  | 0.05 ± 0.00 |  | 0.08 ± 0.00 |  |
| Col20140090 | *E. coli* | TEM-1, CTX-M-15, OXA-1, OXA-1-like | 0.05 ± 0.00 |  | 0.06 ± 0.00 |  | 0.09 ± 0.00 |  |
| Abs<sup>450</sup> positive cut-off<sup>d</sup> |  |  | 0.08 |  | 0.08 |  | 0.09 |  |

<sup>a</sup>RUBLA isolates (20) belong to ARSIA (Association Régionale de Santé et d'Identification Animale), Ciney, Belgium. Col, PEP and CNR isolates belong to the National Reference Center for Antimicrobial Resistance in Gram -, CHU UCL Namur (Mont-Godinne), Belgium. <sup>b</sup>Gene content of RUBLA isolates was determined by Whole Genome Sequencing (WGS) while gene content of Col, PEP and CNR isolates was determined by PCR and amplified fragment sequencing. <sup>c</sup>MutAmpC: chromosomal overexpressed AmpC presenting three mutations in the promotor at positions -1, -18 and -42. <sup>d</sup>The
Abs<sup>450</sup> positive cut-off values were calculated as an average Abs<sup>450</sup> value of the strain *E. coli* DH5α presenting no bla genes plus three times the standard deviation.

### Effects of the VHHs on the enzymatic activity of CMY-2: kinetic characterization

The CMY-2 activity for 4 β-lactam substrates was studied in the presence of the VHH cAb_CMY-2_(254) (Fig. 8). The 4 substrates were chosen based mainly on nature of the side chain of carbon C2 for penicillin and the C3 for the three cephalosporins (Fig. S2). This selection aims to potentially provide a link between the nature of the side chains of these antibiotics and the strength of inhibition. The results highlight that the residual activity of cAb_CMY-2_(254)/CMY-2 complexes, at the higher molar ratio tested, against the three cephalosporins was comprised between 10 and 15 % compared to the activity of the free enzyme. On the other hand, the residual activity for the ampicillin hydrolysis plateaued at 40 %, indicating an inhibition of CMY-2 activity for this substrate less efficient and probably following another mechanism of inhibition. Finally, a similar inhibitory profile was observed for CMY-2 in complex with VHHs cAb_CMY-2_(250) and cAb_CMY-2_(272) for the hydrolysis of nitrocefin (Fig. S3). These results are in full agreement with those of competitive binding assessments (Fig. 4) and support the hypothesis that the three VHHs bind to an overlapping epitope.

**FIG 8.**
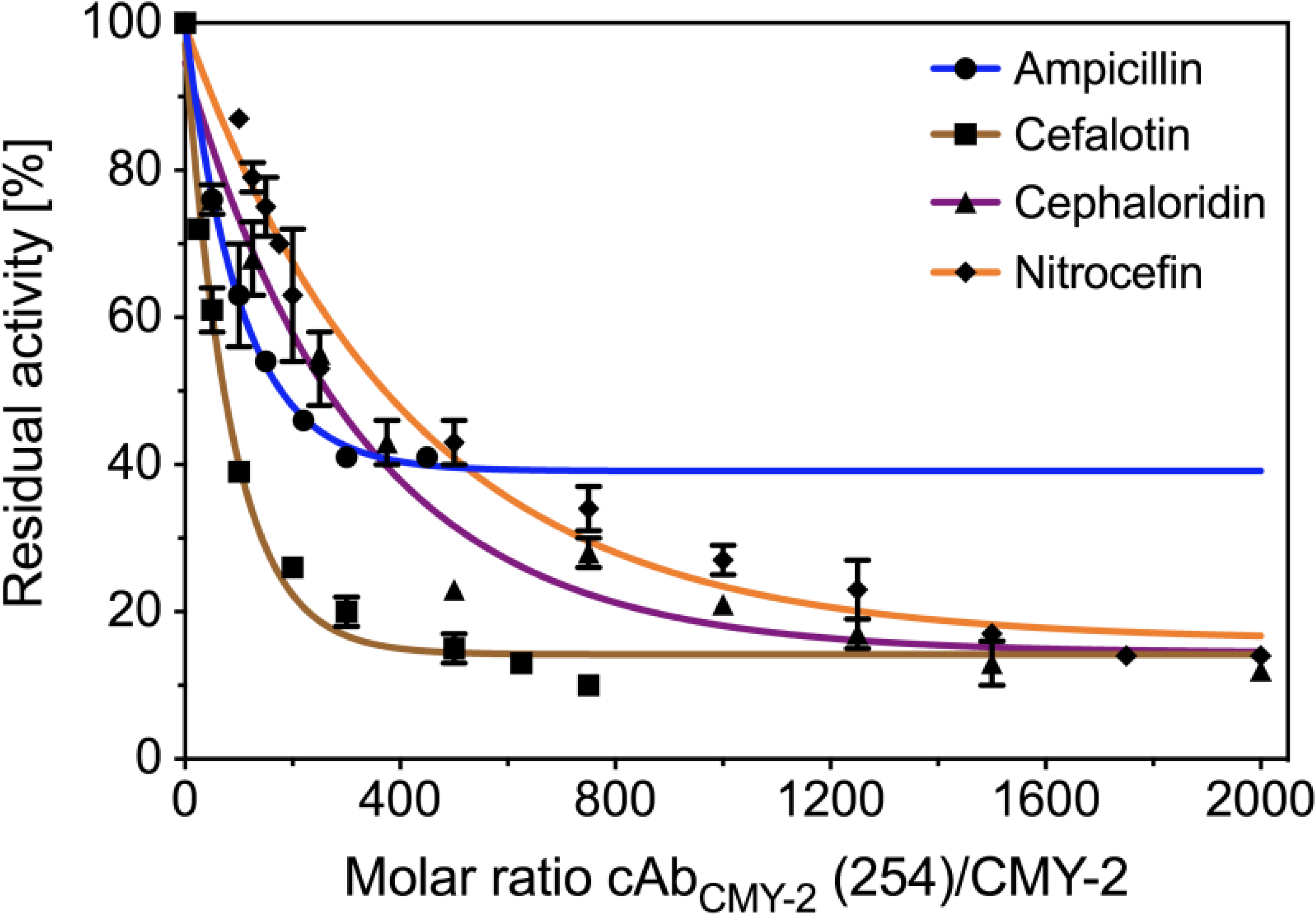
Residual activity of CMY-2 in complex with cAb_CMY-2_(254) for β-lactam ring substrates. Substrates corresponded to ampicillin (blue line), cefalotin (brown line) and cephaloridin (purple line) at 100 μM and nitrocefin (orange line) at 40 μM. Concentrations of CMY-2 used for each substrate described above were 5 nM, 1 nM, 0.2 nM and 0.5 nM, respectively. All data were fitted on a one phase exponential decay equation from graph prism program with values resulted from two experiments realized independently.

Linearization of the substrate hydrolysis curves were carried out from the equation II (TEXT. S1), in order to derive the steady-state kinetic parameters of CMY-2 alone or in interaction with VHHs. The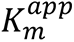 values of CMY-2 for the nitrocefin was not affected by the presence of the VHH, while the deacylation constant 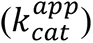 decreased in presence of increasing concentration of cAb_CMY-2_(254) (Fig. 9A). These observations suggest a non-competitive inhibition trend where the VHH did not prevent the interaction between the substrate and the active site of CMY-2. Nevertheless, this VHH could affect the stability of the acyl-enzyme and/or the deacylation phenomenon. Moreover, the plot illustrating 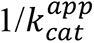 in function of inhibitors concentration (Fig. 9B) displayed a linear trend suggesting a pure non-competitive inhibition. Therefore, the ESI complex was poorly active when the concentration of the VHH cAb_CMY-2_(254) was significantly higher than the inhibition constant value (K_i_). Based on this model, the values of the theoretical parameters α and β were estimated at 1 and 0, respectively (Scheme 1, Material and Methods, steady-state kinetics studies). The K_i_ of the VHH cAb_CMY-2_(254) for CMY-2 was determined from the equation IV (TEXT. S1) and is equal to 88 ± 3 nM which is in good agreement with the equilibrium constant of dissociation of the complex assessed by BLI (TABLE 1).

**SCHEME 1.**
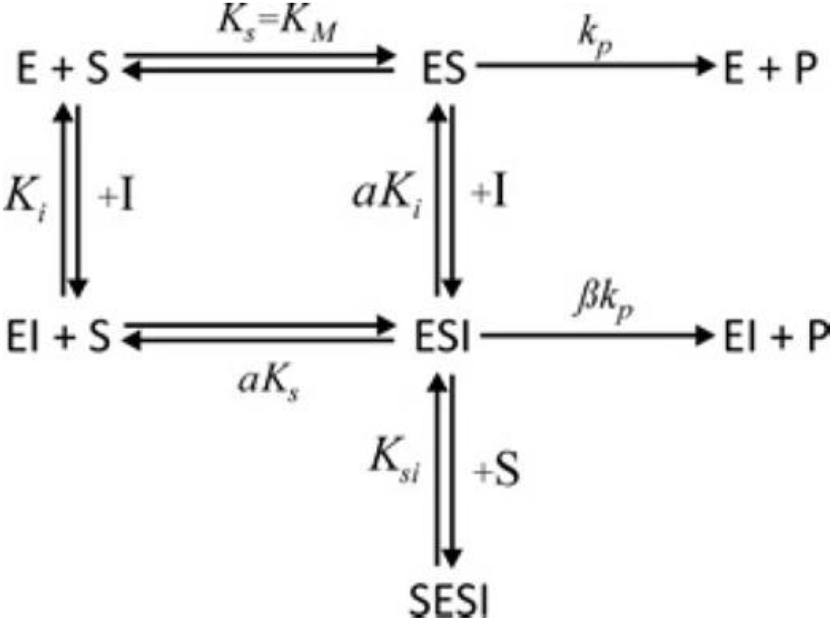
General kinetic model

**FIG 9.**
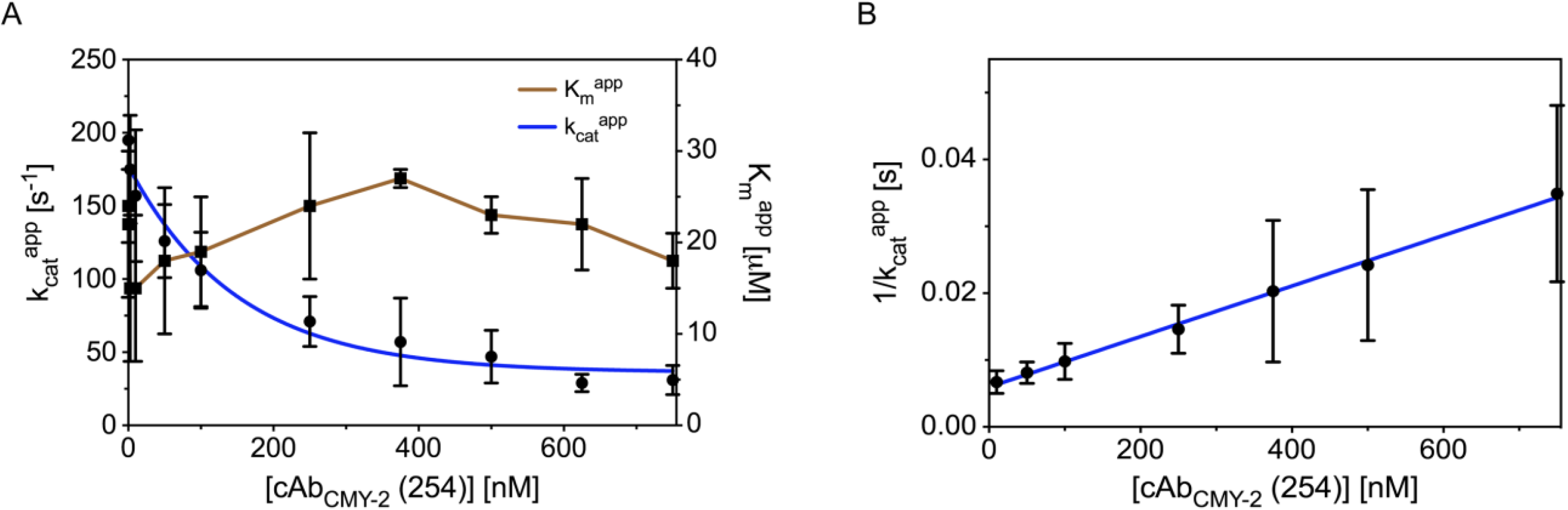
Inhibitory model of CMY-2 activity for the nitrocefin by the VHH cAb_CMY-2_(254). (A) 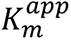 (blue line) and 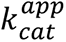 (brown line) parameters derived from the complete hydrolysis of 40 μM of nitrocefin by CMY-2 in complex with the VHH cAb_CMY-2_(254). Experiments were performed using CMY-2 at a concentration of 0.5 nM. 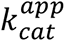 data were fitted using the one phase exponential decay equation from graph prism. (B) Trend of 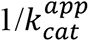 values in function of VHH cAb_CMY-2_(254) concentration. All values resulted from three independent experiments.

Residual activities measurements suggested a similar inhibition pattern of CMY-2 activity by the VHH cAb_CMY-2_(254) for the hydrolysis of the three tested cephalosporins (Fig. 8). Thereby, considering 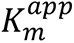 unchanged for any concentrations in inhibitors, the VHH also behaved as a pure non-competitive inhibitor (Fig. 10 A & B) with K_i_ values of 48 ± 10 nM and 107 ± 13 nM for the cephaloridin and the cefalotin, respectively. On the contrary, for ampicillin (Fig. 10C), the hyperbole tendency of 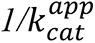 as function of VHH concentration was indicative of a mixed non-competitive inhibition. In this case, the parameter β was equal to 0.41 ± 0.01 and the K_i_ value equals to 352 ± 62 nM (equation V, TEXT. S1). Our data indicated that, for ampicillin, the ESI complex presented a reduced but not abolished activity compared to the ES complex when the concentration of cAb_CMY-2_(254) is significantly higher than the inhibition constant value (K_i_).

**FIG 10.**
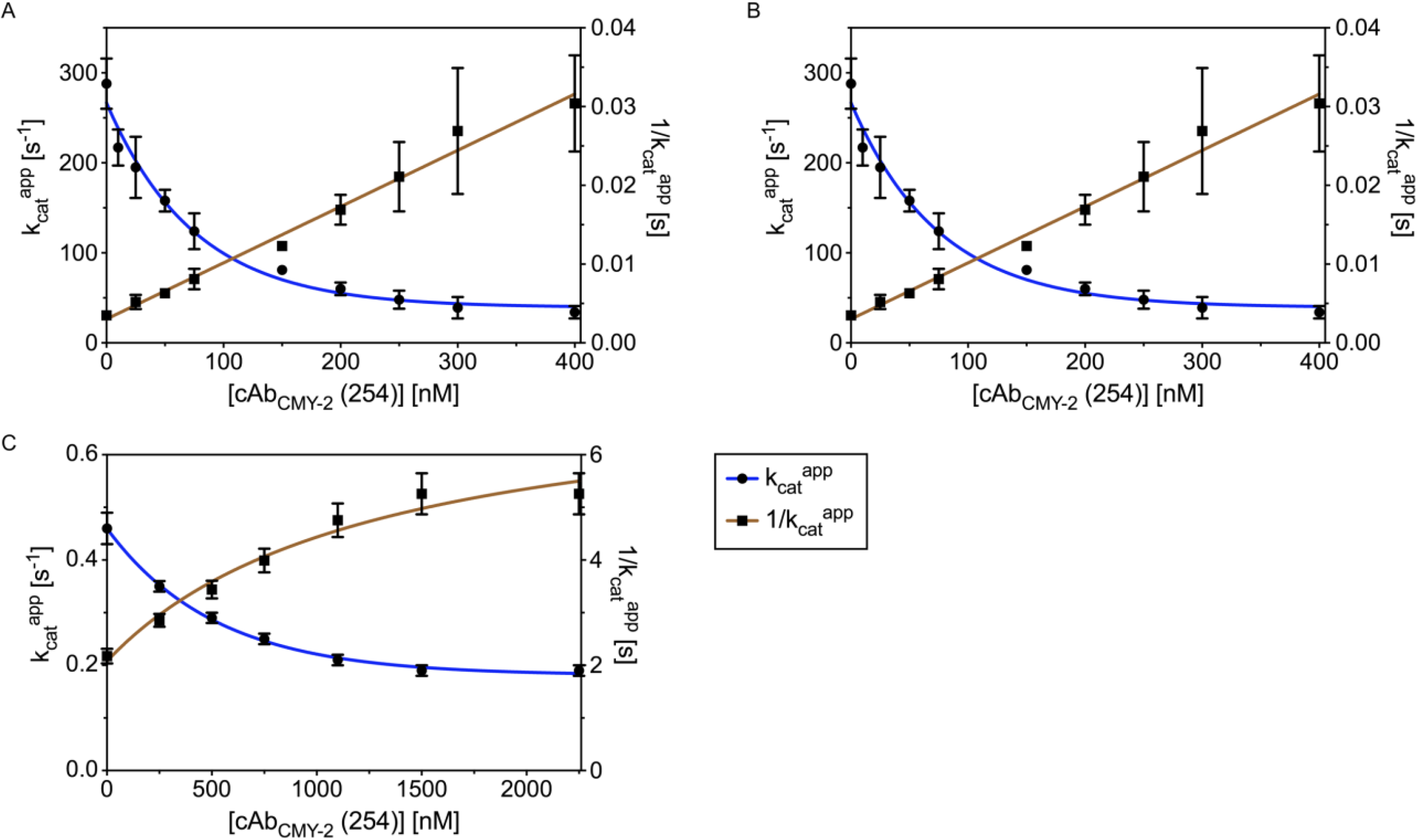
Inhibitory model for CMY-2 activity for the cephaloridin (A), the cefalotin (B) and the ampicillin (C) by cAb_CMY-2_(254). 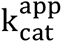 (blue) and 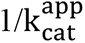 (brown) parameters were obtained from the linear phase (equation III, TEXT. S1) of hydrolysis of the corresponding substrate. Concentrations of CMY-2 were at 0.2 nM, 1 nM and 5 nM for cephaloridin, cefalotin and ampicillin, respectively.

### Structural characterization of the cAb_CMY-2_(254)/CMY-2 complex

The crystal of the cAb_CMY-2_(254)/CMY-2 complex belonged to the P6_2_22 space group and diffracted at a resolution of 3.2 Å. All data and refinement statistics are summarized in Table 3. The asymmetric unit contained one complex CMY-2/VHH. The model includes residues K3 to Q361 in the β-lactamase molecules and residues Q1 to H124 in the cAb_CMY-2_(254) with the exception of residues G108 and E109 in the CDR3 which were not defined in the electronic density.

**Table 3.** Data collection and refinement statistics.

| **Crystal** | cAb<sub>CMY-2</sub>(254)/CMY-2 |
| --- | --- |
| PDB code | 7PA5 |
| **Data collection** |  |
| Space group | P 6<sub>2</sub>22 |
| Cell constants |  |
| a, b, c [Å] | 95.12, 95.12, 242.94 |
| α, β, γ [°] | 90.00, 90.00, 120.00 |
| Resolution range [Å]<sup>a</sup> | 48.88 – 3.18 (3.37-3.18) |
| Rmerge [%]<sup>a</sup> | 71 (342) |
| <I>/<σI><sup>a</sup> | 7.1 (1.28) |
| Completeness [%]<sup>a</sup> | 99.6 (97.9) |
| Redundancy<sup>a</sup> | 37.8 (34.8) |
| CC (1/2)<sup>a</sup> | 0.999 (42.9) |
| **Refinement** |  |
| No. of unique reflections | 11518 |
| R work [%] | 0.2067 |
| R free [%] | 0.2502 |
| No. atoms | 7475 |
| Protein | 7467 |
| Solvent | 8 |
| RMS deviations from |  |
| Bond lengths [Å] | 0.004 |
| Bond angles [°] | 1.03 |
| Mean B factor [Å<sup>2</sup>] | 78.0 |
| Ramachandran plot: |  |
| Favored region [%] | 96 |
| Allowed regions [%] | 4 |

<sup>a</sup>Values in parentheses are related to high resolution shell.

The binding area between the two proteins was about 950 Å^2^. The VHH cAb_CMY-2_(254) interacted via their CDR1 and CDR3 loops at junction between the α and the α/β domains of CMY-2 (Fig. 11 A & B). In more details, a first hydrophobic cluster was formed by residues V2 (N-terminal end), residues F27 and Y32 from CDR1 and I98 from CDR3 of the VHH cAb_CMY-2_(254) and residues K290, V291, A294 and L296 located on the helix α12 and the β-strand β13 of CMY-2 (Fig. 11C).

**FIG 11.**
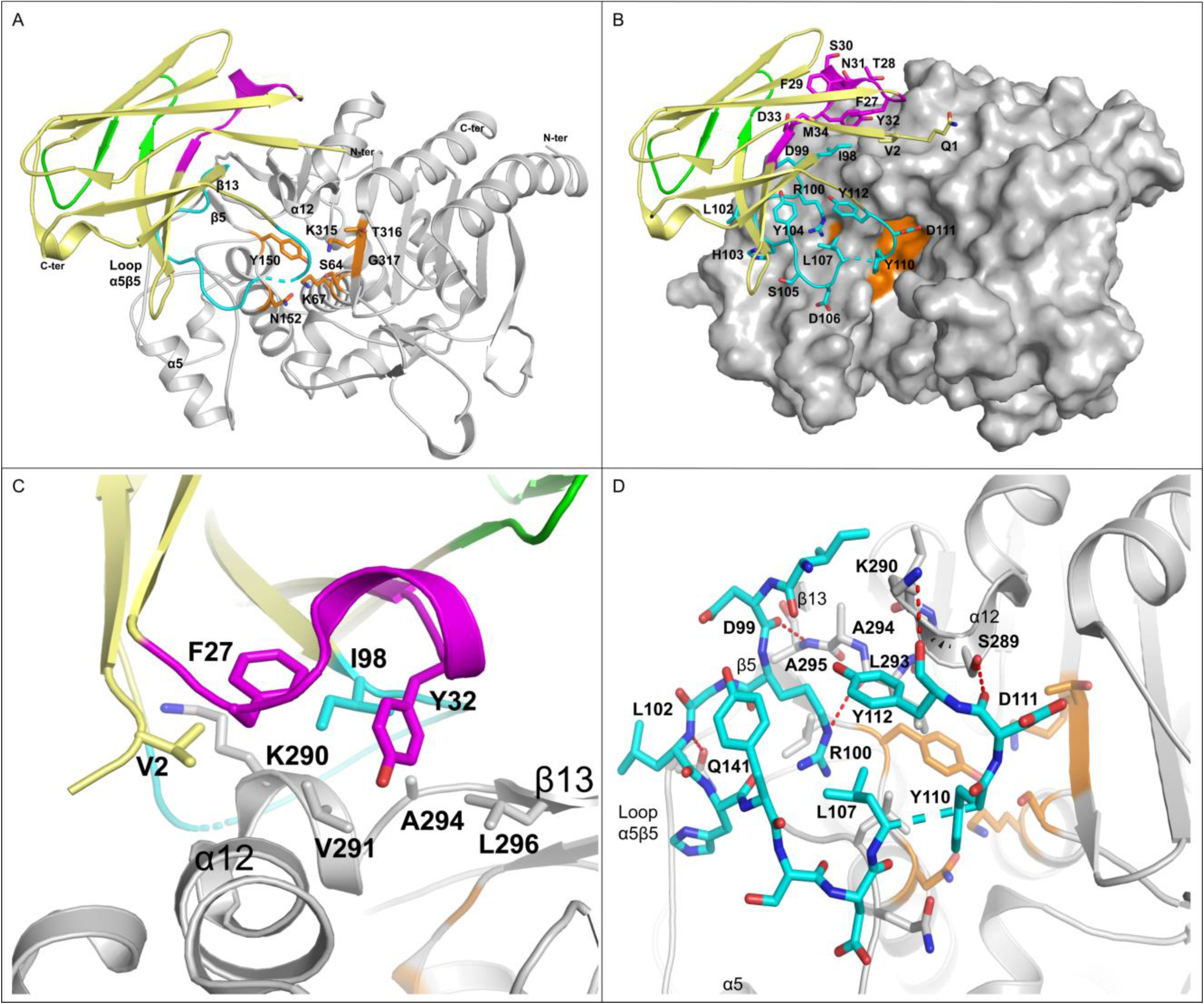
Binding molecular characterization of the complex cAb_CMY-2_(254)/CMY-2. (A) Cartoon representing the overall view of the complex cAb_CMY-2_(254)/CMY-2. (B) Surface representation of CMY-2 in the complex cAb_CMY-2_(254)/CMY-2. (C) Hydrophobic interactions between the CDR1 and the N-terminal extremity of the VHH and CMY-2. (D) Hydrogen bonds (H-bonds) between the CDR3 of the VHH and CMY-2. CDR1, CDR2 and CDR3 of the VHH are colored in purple, green and cyan, respectively, while frameworks are represented in yellow. CMY-2 is representing in grey while residues constituting the motif 1 (S_64_XXK_67_), the motif 2 (Y_150_XN_152_) and the motif 3 (K_315_T_316_G_317_) of the CMY-2 active site are colored in orange. Hydrogen bonds are highlighted by a red dotted line. Residues G108 and D109 from the CDR3 of the VHH cAb_CMY-2_(254) are not illustrated in the model due to a lack of information in the electronic density.

Moreover, the N-terminal residues of the CDR3 (i.e. D99, R100 and L102) established H-bonds with the main chain of residues L293 and A295 located respectively on the helix α12 and the β-strand β13 and with the residue Q141 found on the α5β5 loop (Fig. 11D). Finally, the residues D111 and Y112 from the C-terminal end of the CDR3 made H-bonds with CMY-2 residues S289 and K290, respectively.

All these interactions resulted in a partially entry of the CDR3 into CMY-2 active site, mediated mainly by the residue Y110 of the VHH (Fig. 11D). However, both residues G108 and E109 were not defined in the electron density, highlighting an important flexibility of this CDR3 region and the VHH inability to enable the entrance of the substrate into the CMY-2 active site.

## Discussion

### Overlapping epitopes of the VHHs

The immunization of alpacas allowed the selection of three VHHs. Competition binding assays highlighted that the three VHH bind to an overlapping epitope on CMY-2. The structure of the complex cAb_CMY-2_ (254)/CMY-2 revealed essentially the insertion of the CMY-2 K290 into a pocket on the surface of cAb_CMY-2_ (254) (Fig. 11C) and a second binding area involving most of the CDR3 (Fig. 11D). The residues forming the pocket are conserved between the three VHHs except for the I98 which is mutated into an alanine (Fig. 1). Despite the CDR3 constitutes the least conserved region among the VHHs, it is probable that the three VHHs share a similar binding mode with different affinities related to the ability of the CDR3 to bind to CMY-2.

### Biochemical features of the VHHs for the CMY-2 sub-family

*In vitro* binding assays demonstrated that cAb_CMY-2_ (254) and cAb_CMY-2_ (272) present a higher specificity for β-lactamases belonging to the CMY-2 sub-family (no recognition of CMY-1 and P99) than cAb_CMY-2_ (250) which also binds P99. The both higher affinity and specificity of the VHH cAb_CMY-2_ (254) justified its use for the screening of bovine and human bacterial isolates by a sandwich ELISA assay. *In vivo* binding assays on bacterial isolates highlighted the ability to detect different variants from the CMY-2 sub-family as CMY-16, −42, −58, −60 and −61. In fact, the sequence implied in the interaction with the VHH is conserved for all variants from the CMY-2 sub-family meaning the high probability to detect also other CMY-2-like β-lactamases. Moreover, we were not able to detect other class C β-lactamases such as ACT-1, DHA-1 and CMY-10. ACT-1 and P99 present a high sequence identity with CMY-2 (Fig S4) providing a similar conformation of the helix α12 (Fig. 12A). However, the Ala295 is substituted by a proline in P99 and ACT-1 what may explain the inability to interact with the VHH. The presence of a proline introduces a steric hindrance in the helix and may displace the H-bonds network stabilizing the VHH/CMY-2 complex. The steric hindrance with the glutamate E294 and the total conformation change of the helix α12 due to a low sequence identity with CMY-2 could prevent the interaction of CMY-1 and CMY-10 with the VHH (Fig. 4.12B).

**FIG 12.**
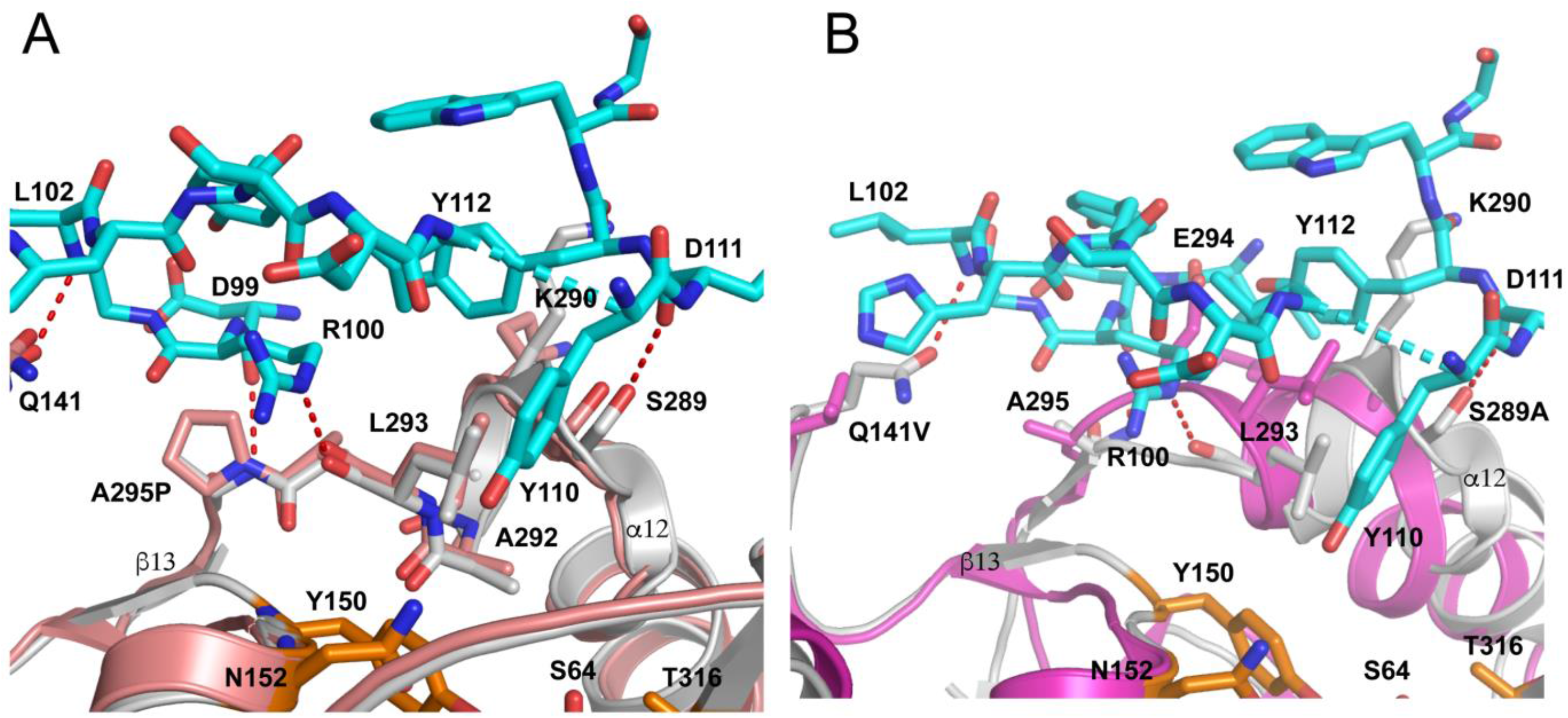
Superposition of complex cAb_CMY-2_ (254)/CMY-2 and P99 (PDB code 1XX2) (A) and CMY-10 (PDB code 1ZKJ) (B). CMY-2, P99 and CMY-10 are colored in grey, red and magenta. Only the CDR3 of the VHH is illustrated in cyan and active site residues in orange. H-bonds are represented by a red dashed line.

### Biochemical features of the polyclonal antibodies against CMY-2

The overlapping epitope on CMY-2 and shared by the three VHHs required the development of rabbit polyclonal antibodies. These antibodies were less specific since they were able to recognize P99 and CMY-1. They correspond to a mix of antibodies probably able to bind to epitopes shared by a large panel of AmpC β-lactamases what could explain their lack of specificity (32). Fortunately, the use of the VHH cAb_CMY-2_ (254) permitted to offset the low specificity of the pAbs for the detection of CMY-2 in the sandwich ELISA.

### Development of tandem-repeats VHH cAb_CMY-2_ (254)_BIV_

Another interesting aspect with the VHHs is the possibility to fuse them in order to decrease the dissociation rate by an avidity phenomenon leading to more stable complexes Antigen/Antibody (35-37). Associated with the multi-avidity ensured by the polyclonal antibodies, this allowed to detect lower quantities of CMY-2 than the monovalent counterpart.

### Applicability in an ELISA

This study presented as more interest the use of the VHH as antibody for the detection of a bête-lactamase. On contrary with monoclonal antibodies, they are easier to produce and purify and present some biochemical features allowing a better stability and solubility. Moreover, they generally display a high affinity and specificity for its antigen essential to obtain the more suitable detection assay as already demonstrated for cancer biomarkers (38, 39). The possible lower affinity of some VHHs can be compensated by the engineering of in tandem-repeats VHHs improving the sensitivity of a detection assay due to an avidity phenomenon.

One goal of the project RU-BLA-ESBL-CPE consisted to develop a sandwich ELISA “type” for the detection of one of the most spread β-lactamase, CMY-2, in bovines and more largely in the animal world. However, our next aim is to develop an Immunochromatographic Lateral Flow Assay that ensures a detection more rapidly and in an easier manner aiming an interesting alternative for veterinarians to phenotypic methods (40).

Finally, despite this test is probably suitable for CMY-2 detection in animals, it stays less applicable in human medicine where phenotypic assays constitute an unavoidable method for selection of the best antibiotic. However, we could imagine use this kind of set up to interpret more easily difficult phenotypic profiles generally found in MDR strains (27), to distinguish plasmid to chromosomic AmpC (41) and to highlight the involvement of an AmpC in a carbapenemase activity of the strain (28).

### The VHH cAb_CMY-2_ (254), a non-competitive inhibitor of the CMY-2 activity

This work allowed also the selection of VHHs which behave as non-competitive inhibitors. The structure of the complex cAb_CMY-2_ (254)/CMY-2 highlighted an important flexibility of the CDR3 loop located in the active site what does not prevent the entry of the substrate in the active site. Nevertheless, the Tyr100 brought by the CDR3 is situated near to the Gln120 which is considered as a crucial residue involved in the stabilization of the acyl-enzyme by the establishment of H-bonds with the C7 amide carbonyl of the substrate (42). Therefore, despite the Y110 does not directly bind the Gln120 in CMY-2, this may impede the stabilization of the acyl-enzyme (Fig. 13). Moreover, the interaction of the VHH around the active site may perturb the dynamic of the enzyme which is known to be essential for the optimal activity of the enzyme (43).

**FIG 13.**
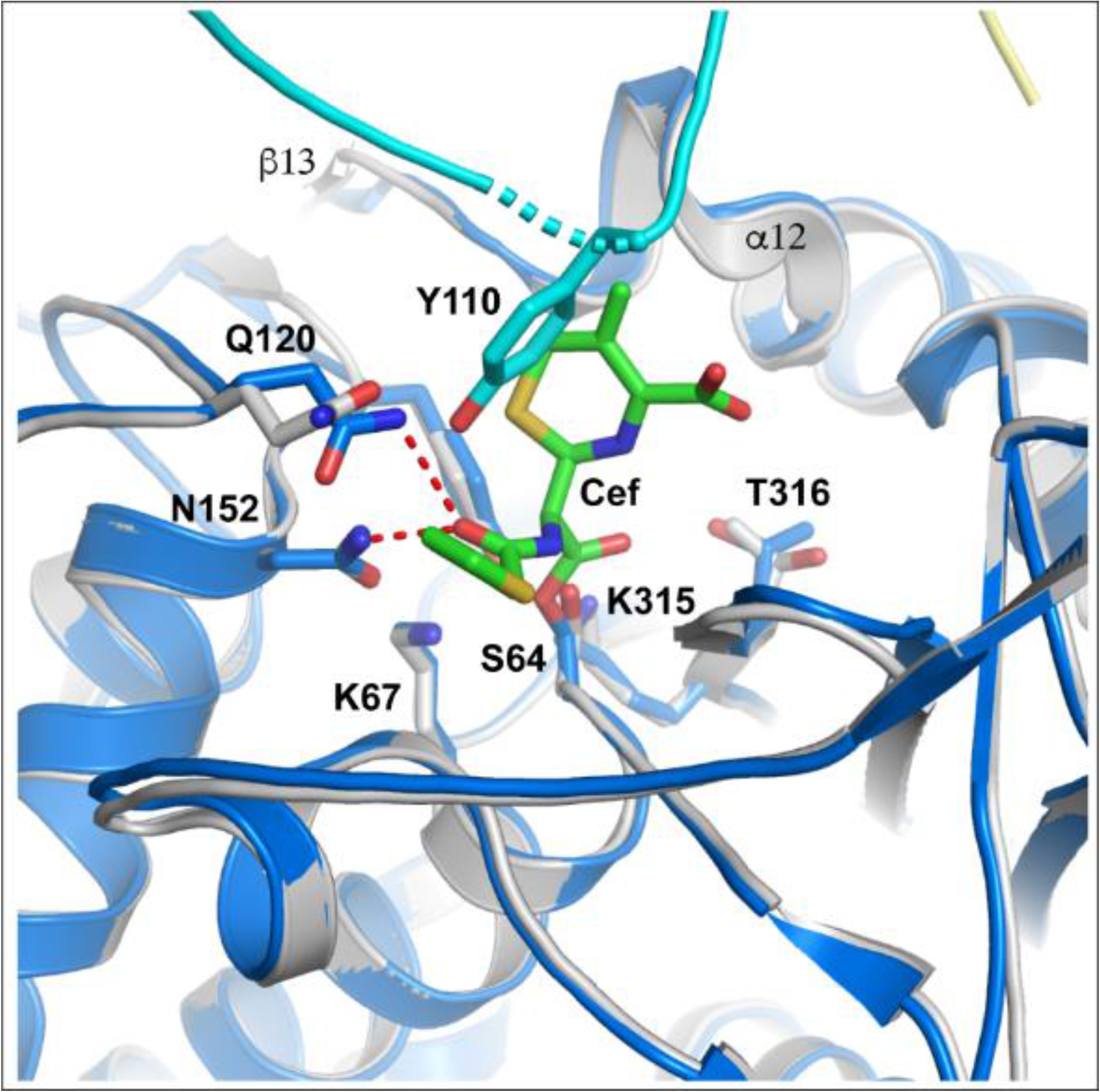
Superposition of the complex cAb_CMY-2_(254)/CMY-2 and the crystal structure of AmpC WT β-lactamase from *E. coli* in complex with a covalently bound cefalotin (PDB code 1KVM). CMY-2 is colored in grey, the AmpC in blue, the cefalotin in green and the CDR3 of the VHH is illustrated in cyan.

Our study provided also the evidence that the mechanism of inhibition can be different in function of the substrate. We found that the VHH behaved as a non-competitive inhibitor with its ability to completely inhibit the activity of CMY-2 for all cephalosporins tested (scheme 1). However, in presence of ampicillin, the complex maintained a reduced activity corresponding to a mixed non-competitive inhibition. The more plausible explanation consists in the fact that the ampicillin can easily diffuse in the active site due to its smaller size resulting in less impact by an eventual steric hindrance and/or motion perturbations.

### Peptidomimetics from VHH, an alternative strategy to classical inhibitors

Despite the VHH is smaller than classical antibodies (15 KDa versus 150 KDa), it stays too bulky to penetrate into the periplasm space of the bacteria. Actually, one strategy in view to minimize the size of inhibitors consists in the development of small peptides by peptidomimetics. These present several advantages: (I) an easier production in large scale-up with a cheaper cost, (II) the low tissues penetration of large molecules and antibodies rendering less efficient the drug delivery and its action and (III) the humanization of therapeutics antibodies which can be laborious and which can lead finally to the development of human anti-mouse antibody (HAMA) (44, 45).

One category implied the development of peptides based on the therapeutics monoclonal antibodies as from the rhuMAb 4D5 (trastuzumab) used in the treatment of the breast cancer where a gene HER-2 is upregulated and induces the cellular proliferation (46). More interestingly, VHHs were also used as scaffold for the development of peptides as against the VEGF (Vascular Endothelial Growth Factor) factor implied in angiogenesis in tumor development (47) or against the receptor β-2 adrenergic associated with chronic inflammation (48).

The main drawback to consider in peptidomimetics consists in generally lower affinities compared to the corresponding antibodies. In fact, we could reach K_D_ values near or upper than 1 μM. However, some studies demonstrated that lower affinities resulted from a decrease of the association constant independently of the dissociation rate (49, 50).

To conclude, peptides remind an important alternative to nanobodies as therapeutic agents due to their smaller size and their interesting pharmaceutical features against some domains as cancers or inflammation. We could consider this type of development against the β-lactamases as CMY-2 which stays more interesting for the veterinarians, but also against more interesting enzymes such as metallo-β-lactamases.

## Material and methods

### Production of the β-lactamase CMY-2

CMY-2 was produced as previously described by *Cedric Bauvois et al., 2005* (21). The CMY-2 protein was stored at −20°C in 50 mM MOPS buffer at pH 7.0 containing 10 % glycerol (w/v) and at −20 °C. Its integrity was verified by Coomassie-stained SDS-PAGE and mass-spectrometry (ESI-Q-TOF). The concentration of the purified enzyme was determined by its absorbance at 280 nm (ε^280^ = 93850 M^-1^ cm^-1^).

### Selection of VHHs by phage display

One alpaca (*V. pacos*) was immunized by six weekly sub-cutaneous injections of 100μg of LPS-free CMY-2 mixed with Gerbu adjuvant. The immune library was constructed following a previously developed protocol from *Conrath et al* (18) while the VHHs selection by phage display and the screening of the selected VHHs were performed as described in *Pardon et al* (51). All details concerning those experiments are described in the supplemental material (TEXT S1).

### Cloning of VHHs genes into pHEN14, scale-up production and purification

Genes coding for VHHs selected by phage display were subcloned into the expression vector pHEN14 between the restriction enzyme sites HindIII in the 5’-extremity and BstEII in the 3’-extremity. This vector derived from the phagemid pHEN6 where the resistance to ampicillin is replaced by the resistance to chloramphenicol and where there is no myc tag (19). Genes coding for the bivalent VHHs, corresponding to two identical VHHs in tandem repeats joined by a peptide linker (GGGS)_3_, were ordered into the pHEN14 from Genecust (Boynes, France) (34). Production of monovalent and bivalent VHHs started with the transformation of competent *E. coli* WK6 with plasmid constructs by thermic shock. Then, the cells were plated on LB agar containing chloramphenicol (25 μg/mL) for selection. VHHs were produced in flasks in a Terrific Broth Medium supplemented by the antibiotic (25 μg/mL) and where a preculture of one colony was added to attempt an initial OD^600^ = 0.2. The growth was performed at 37°C until an OD^600^ ≍ 0.8 before addition of 1 mM IPTG to induce the production of the VHHs overnight at 28°C. The cells were harvested and a periplasmic extraction by osmotic-shock was carried out with a solution containing 0.5 M sucrose. This extraction was followed by an affinity chromatography with an HisTrap HP Ni-nitrilotriacetic acid column (Cytiva) and a purification by size-exclusion chromatography (Superdex75). Purified VHHs were conserved in a 50 mM PBS pH 6.1. The purity and the integrity of the VHHs were verified by Coomassie-stained SDS-PAGE and masse spectroscopy (ESI-Q-TOF).

### Immunization of rabbits and purification of polyclonal antibodies (pAbs)

Polyclonal antibodies (pAbs) were obtained by rabbit immunization realized by the CER Group (Marloie, Belgium), that consisted in four injections of 500 μg of CMY-2 all two weeks in a standard subcutaneous way. Then, sera recovered from blood were conditioned in a 50 mM PBS pH 7.4 buffer and pAbs were purified with a HiTrap Protein A HP antibody purification column (Cytiva) where the elution buffer corresponded to 20 mM Glycine pH 2.0 buffer. Fractions containing the pAbs were pooled and dialyzed against a 50 mM PBS pH 7.4 buffer overnight at 4°C. The integrity and purity of the pAbs were assessed by Coomassie-stained SDS-PAGE while concentration of the pAbs was measured by Bicinchoninic Acid Assay (BCA).

### In-vitro biotinylation of antigen and antibodies

Bio-layer interferometry experiments (BLI) and ELISAs tests may require biotinylated proteins. To this aim, we used the EZ-link^@^NHS-PEG_4_ biotin kit (ThermoScientific, United States) to covalently bind biotin molecules on lysine residues of proteins. The chemical reaction was performed at room temperature, for 30 minutes and with a [Biotin]:[protein] ratio of 3:1. The excess of biotin was removed by the elution of the reaction mixture on Sephadex G25 column. The labelled protein was conserved in 50 mM PBS at pH 7.5 at a final concentration between 100 and 500 μg/mL.

### Kinetic characterization by bio-layer interferometry

All bio-layer interferometry (BLI) experiments were performed on the OCTET HTX instrument (ForteBio, Sartorius) at 30°C using 96-well black polypropylene microplates (Greiner BioOne, Belgium). All proteins were diluted in a kinetic buffer (50 mM PBS pH 7.4 supplemented with 0.1 % BSA (w/v) and 0.05 % tween-20 (v/v). All data were analyzed by the Octet software version 12.0 (Sartorius, France).

The specificity of the binding consisted in the immobilization of 2 μg/mL of purified VHHs on Anti-His coated sensors (His1K, Sartorius) via their His_6_ tag. Then, a baseline was monitored with the kinetic buffer for 60 s. Binding to the VHHs coated on the sensor was monitored by incubating, for 120 s, the VHH in presence of a solution of 500 nM of antigens representing all classes of β-lactamases: TEM-1 for class A, VIM-4 for class B, CMY-1, CMY-2 and P99 for class C and OXA-48 for class D. The dissociation kinetic constant of the complex was monitored for 300 s by incubating the sensor in the kinetic buffer. Moreover, for the quantitative binding assays, the conditions for each VHH were as follows: i) cAb_CMY-2_ (250) was assessed on 10 and 60 s of association and dissociation, respectively, using a range of CMY-2 concentration between 50-250 nM; ii) cAb_CMY-2_ (254) on 60 and 600 s and using a range between 40 and 450 nM; iii) cAb_CMY-2_ (272) on 30 and 180 s and using a range between 20 and 110 nM. Kinetics constants (k_on_ and k_off_) and equilibrium constant (K_D_) were calculated using a 1:1 interaction model with a global fit based on at least seven analyte concentrations indicated on all sensorgrams.

Avidity studies were achieved using streptavidin bio-sensors (SA sensor, Sartorius) where biotinylated CMY-2 was immobilized for 30 to 50 minutes at a concentration comprised between 10 and 50 μg/mL. A quench reaction was realized by incubating biocytine 10 μM for 300 s. Then, the binding of the monovalent and the bivalent VHHs was monitored for 30 s using a range of CMY-2 concentration between 150-1000 nM and 75-375 nM, respectively. The dissociation of the complexes was measured for 600 s in the kinetic buffer. The binding of the rabbit polyclonal antibodies (pAbs) to the antigen was monitored for 60s (12.5-200 nM of CMY-2) and the dissociation of the complexes was measured for 600 s. A global fit based on 5 analyte concentrations was realized only for dissociation constant (k_off_) thanks to an exponential decay mathematic model. Specificity binding of the pAbs was undertaken following the same setup except the association that was measured for 300 s using 500 nM of pAbs.

Competition binding assays were performed by a premix method with streptavidin bio-sensor (SA sensor, Sartorius). Firstly, a biotinylated VHH (2 μg/mL) was immobilized on the sensor to reach a variation of the signal (Δλ) of around 1 nm. Complexes VHH/CMY-2 were obtained by incubating CMY-2 (200 nM) and the VHH (100 nM-4 μM) for 15 minutes at 30°C. Then, the solutions were loaded in order to assess the association between the immobilized VHH and the complexes VHH/CMY-2. Binding rates were measured for the first 120 s of the association phase with an exponential mathematics model.

### pAbs specificity by indirect ELISA

The specificity of pAbs directed against CMY-2 (Anti-CMY-2 pAbs) was determined by an indirect ELISA. To this aim, 500 ng of antigens representing all classes of β-lactamases and diluted in 50 mM MES pH 5.5 buffer were immobilized by absorption on a 96-well NUNC maxisorp (ThermoScientific, United States) overnight at 4°C. All non-specific sites were saturated using 1 % BSA (w/v) for two hours. Then, 500 ng of anti-CMY-2 pAbs were added in each well. The assay was revealed by a 1/2000 diluted goat anti-rabbit antibody conjugated to horseradish peroxidase (HRP) (Abcam, UK). All steps were followed by 5 washes with 50 mM pH 7.5 buffer with 0.05 % tween-20. Antibodies were diluted in the washing buffer and all incubations were performed for 1 hour at 28°C. TMB (*3,3’,5,5’-Tetramethylbenzidine*, Merck, Germany) substrate was used for system revelation. The reaction was quenched with 1M H_3_PO_4_ and the plates were read at 450 nm using an Infinite M200 Pro microplate reader (Tecan, Switzerland).

### Sandwich ELISA assay development for CMY-2 detection

A sandwich ELISA for CMY-2 detection was designed to investigate the limit of detection (LOD) and the specificity of the different assays formats. To this aim, several combinations of capture and detection VHHs and anti-CMY-2 pAbs were tested. Briefly, 500 ng of biotinylated VHH cAb_CMY-2_ (254) (monovalent or bivalent) or 2 μg of biotinylated anti-CMY-2 pAbs were used as capture agent on a 96-well NUNC streptavidin polysorb plate incubated overnight at 4°C. The plate was blocked by a 1 % (w/v) BSA solution. Then, the purified CMY-2 was added in serial dilutions from 10^-4^ to 2 μg/mL to determine the LODs of the four assay combinations. The LODs values were calculated with a sigmoidal model on graph prism (equation I, TEXT S1).

The specificity of the assays was evaluated with 200 ng of the 7 β-lactamases belonging to the four classes of β-lactamases. At least three wells where antigen was omitted were used as blank. Additionally, the detection of CMY-2 produced by human and bovine bacterial isolates was performed as follow. The different strains were grown in TB medium supplemented by 100 μg/mL ampicillin for 4 hours at 37°C. Strains were lysed by sonication with a Bioruptor Plus (Diagenode, Belgium) and centrifuged at 18000g in order to recover the bacterial content. An *E. coli* DH5α strain was used as negative control. All detection assays on bacterial isolates were realized by using 5 μg of bacterial crude extract. Detection of CMY-2 was performed by adding 500 ng/well of pAbs themselves followed by the addition of a 1/2000 diluted goat anti-rabbit antibody conjugated to HRP (Abcam, UK) or by adding 200 ng/well of monovalent or bivalent cAb_CMY-2_ (254) recognized by 1/2000 diluted rabbit anti-HCAbs antibody conjugated to HRP (Genscript, United States). TMB was used as substrate while reaction was stopped by 1 M H_3_PO_4_. Abs^450^ was recorded using an Infinite M200 Pro microplate reader (Tecan, Switzerland). All steps described above were performed for one hour at 28°C and were followed by 5 washes of 50 mM PBS pH 7.5 buffer where 0.05 % tween-20 was added.

### Steady-state enzymatic kinetics

Steady-state enzymatic kinetics were performed at 30°C using a 50 mM PBS pH 7.5 supplemented with 50 μg/mL BSA. Absorbances were measured with a Specord 75 spectrophotometer (AnalytikJena, Germany) and a SpectraMx M2 microplate reader (Molecular Devices, United States). Initial rates and complete hydrolysis of substrate were measured for the hydrolysis of : 100 μM ampicillin (Δε^235^ = −820 M^-1^ cm^-1^), 100 μM cefalotin (Δε^273^ = −6300 M^-1^ cm^-1^), 100 μM cephaloridin (Δε^260^ = −10000 M^-1^ cm^-1^) and 40 μM nitrocefin (Δε^482^ = + 15000 M^-1^ cm^-1^). The enzyme CMY-2 concentration used to hydrolyze the various substrates was comprised between 0.2 nM to 5 nM and was mixed with increasing amounts of the VHH cAb_CMY-2_ (254) (0-1500 nM). All steady-state kinetics constants were measured by using equations described in the supplemental material (TEXT S1).

The kinetic model for the inhibition events of CMY-2 activity by the VHHs is described in scheme 1 (19) where K_i_ corresponds to the dissociation constant of the inhibitor. The α parameter is the degree at which the inhibitor influences the affinity of the enzyme for its substrate, while the β parameter is the activity of the tertiary complex ESI compared to the activity of the complex ES. The constant k_p_ corresponds to the turnover rate constant (k_cat_).

### Crystallization conditions

Crystals were grown at 20°C using the sitting drop vapor diffusion method. The drop contained 0.2 μL of CMY-2 in complex with VHH cAb_CMY-2_(254) at a concentration of 14 mg/mL and 0.2 μL of 0.1 M TRIS-HCl pH 8.5 buffer with 1.4M (NH_4_)_2_ tartarate. The crystal was transferred in a cryo-protectant solution containing 50% (v/v) polyethylene glycol 400 and 50 % (v/v) glycerol and frozen in liquid nitrogen.

### Data collection, phasing, model building and refinement

Data were collected at the Proxima 1 beamline of the Soleil synchrotron (Saint Aubin, France). Indexing, integration and scaling of the data were performed using XDS (52). Initial phases were obtained by molecular replacement with the CMY-2 structure (PDB code 1ZC2) and a lama antibody fragment bound to Galectin 10 (PDB code 6GKU, 53) as a search models using Phaser (54). The structure was built with Coot (55) and refined with Phenix refine (56). Figures were prepared using PyMOL (The PyMOL Molecular Graphics System, Version 2.4.1 Enhanced for Mac OS X, Schrödinger, LLC.).

## Acknowledgments

We thank the Protein Factory Platform at University of Liège for providing material necessary for protein purification and to provide some purified β-lactamases necessary for the specificity binding experiments. We would like also to thank the Robotein Platform for the opportunity to use the OCTET HTX robot necessary for all binding characterizations.

This work was supported by the Belgian Federal Public Service Health, Food Chain Safety and Environment [Grant No. RF 17/6317 RU-BLA-ESBL-CPE] and the FNRS [Grant J0081-20-CDR].

